# Regional, layer, and cell-class specific connectivity of the mouse default mode network

**DOI:** 10.1101/2020.05.13.094458

**Authors:** Jennifer D. Whitesell, Adam Liska, Ludovico Coletta, Karla E. Hirokawa, Phillip Bohn, Ali Williford, Peter A. Groblewski, Nile Graddis, Leonard Kuan, Joseph E. Knox, Anh Ho, Wayne Wakeman, Philip R. Nicovich, Thuc Nghi Nguyen, Emma Garren, Cindy T. J. van Velthoven, Olivia Fong, David Feng, Maitham Naeemi, Alex M. Henry, Nick Dee, Kimberly A. Smith, Boaz P. Levi, Lydia Ng, Bosiljka Tasic, Hongkui Zeng, Stefan Mihalas, Alessandro Gozzi, Julie A. Harris

## Abstract

The evolutionarily conserved default mode network (DMN) is characterized by temporally correlated activity between brain regions during resting states. The DMN has emerged as a selectively vulnerable network in multiple disorders, so understanding its anatomical composition will provide fundamental insight into how its function is impacted by disease. Reproducible rodent analogs of the human DMN offer an opportunity to investigate the underlying brain regions and structural connectivity (SC) with high spatial and cell type resolution. Here, we performed systematic analyses using mouse resting state functional magnetic resonance imaging to identify the DMN and whole brain axonal tracing data, co-registered to the 3D Allen Mouse Common Coordinate Framework reference atlas. We identified the specific, predominantly cortical, brain regions comprising the mouse DMN and report preferential SC between these regions. Next, at the cell class level, we report that cortical layer (L) 2/3 neurons in DMN regions project almost exclusively to other DMN regions, whereas L5 neurons project to targets both in and out of the DMN. We then test the hypothesis that in- and out-DMN projection patterns originate from distinct L5 neuron sub-classes using an intersectional viral tracing strategy to label all the axons from neurons defined by a single target. In the ventral retrosplenial cortex, a core DMN region, we found two L5 projection types related to the DMN and mapped them to unique transcriptomically-defined cell types. Together, our results provide a multi-scale description of the anatomical correlates of the mouse DMN.

## Introduction

Large-scale, distributed brain networks support sensory perception, cognition, and motor output. These networks are typically identified by temporally correlated activity (functional connectivity, FC) or by measures of structural connectivity (SC). Some distributed networks, called resting state (rs) networks, are characterized by temporally correlated *intrinsic* activity, but their functions are less well understood. One of several rs networks is the cortical default mode network (DMN), an anatomically distributed set of brain areas characterized through rs functional magnetic resonance imaging (rsfMRI) by low-frequency (<1 Hz) correlated activity in the absence of a goal-directed task or stimulus^1^. Functions attributed to the DMN include mind wandering or spontaneous cognition, mood and emotional processing, and learning and memory^1^. Notably, DMN analogs have been identified in monkeys^2^, rats^3,4^, and mice^5–8^. Conservation of this large-scale network across multiple mammalian species suggests a central role in organizing healthy brain activity. Indeed, the DMN is particularly vulnerable in Alzheimer’s Disease (AD), where pathological changes in network activity measured with rsfMRI correlate with disease stage, ultimately ending in a breakdown of connectivity^9–11^. This selective functional network degeneration is closely associated with the spatial and temporal distribution of other AD-related pathologies, specifically the early accumulation of amyloid deposits^12^. Similarly, the DMN comprises high centrality hub regions^12,13^ that are thought to serve as vulnerability points for several psychiatric disorders such as autism and schizophrenia^14,15^.

The brain regions that comprise the human DMN are predominantly cortical, symmetric across hemispheres, and broadly distributed in anterior and posterior regions; including the medial prefrontal cortex, precuneus, posterior cingulate cortex (PCC), retrosplenial cortex (RSP), and lateral posterior parietal cortex^12,16^. Areas outside the cortex are also sometimes included or considered part of the DMN, *e.g.* entorhinal cortex, hippocampus, and thalamic nuclei^17–19^. The strong FC between these regions during task-free states must be constrained by a foundation of anatomical connectivity patterns. Investigations into the organization of human cortical SC, based on mapping large fiber tracts using diffusion tensor or spectrum imaging (DTI/DSI), led to the identification of a core set of interconnected cortical areas that clearly correspond to posterior parts of the DMN^20^. Significant correlations between SC and FC in human DMN regions are also reported^20–22^. Still, our understanding of the anatomical basis of the DMN is severely limited by the relative coarseness of macroscale connectome mapping in humans.

Higher resolution cortical connectomes based on anterograde or retrograde tracing methods are available for several species in which DMN analogs have been identified, including macaque, rat, and mouse^23–27^. These provide directional and quantitative measurements of SC at the *mesoscale;* a level spanning regions, cell populations, classes and types^28^. Mesoscale connectome analyses reveal conserved rules of cortical connectivity across species, *e.g.,* a unique “fingerprint” of targets reached by each cortical area, strong distance-dependence with a log-normal distribution of connection weights^24,26^, and modular structure with high clustering coefficients and some very highly connected nodes (“hubs”)^23,29–32^. Mesoscale SC at a regional level is generally well correlated with FC^33–35^, and comparisons within the DMN have revealed direct anatomical connections between areas^6,32,34,36,37^. However, the correspondence is not 1:1, unsurprising given that brain regions participate in multiple, partially overlapping functional networks. Network multiplexing may instead be achieved at the level of cell populations, class, or even type using one of the organizational motifs identified by analyses of mesoscale connectivity datasets, such as cell class-specific projection patterns^38^ or laminar-based separation of feedforward and feedback connections^25,39^.

Cross-species conservation of both cortical structural connectivity patterns and rs networks like the DMN suggest that we will gain relevant insight into the anatomical structure of human brain-wide functional networks by studying circuitry in animal models^6,40^. In the mouse, we also have the advantage of higher resolution anatomical descriptions through the rapidly growing census of cell types^41–46^ and a genetic toolbox for accessing and manipulating cell type-specific circuitry^47–50^. These genetic tools allow us to reproducibly identify, access, and even modify the activity of specific cells based on their transcriptomic, morphological, or projection types, and to more confidently compare data across species, individuals, and modalities.

Here, we provide a detailed spatial description of the brain regions that comprise the mouse DMN at 100 μm voxel-level resolution by registering a rsfMRI DMN mask^51^ with the fully annotated 3D Allen Mouse Common Coordinate Framework Reference Atlas (CCFv3^52^). By examining the relationship between the DMN and structural connectivity using anterograde axonal tracing experiments from the Allen Mouse Brain Connectivity Atlas^24,25^, we show that voxels and regions within the DMN are strongly and preferentially connected to other regions in the DMN. We report that the DMN overlaps with two structural network modules in anterior and posterior locations (prefrontal and medial modules). Furthermore, we find that axonal projections arising from distinct layers and projection neuron classes relate in different ways to the DMN mask. Specifically, the vast majority of projections from layer (L) 2/3 neurons in DMN regions target other DMN regions, whereas L5 neurons target areas both in and out of the DMN, particularly in the posterior DMN region, ventral retrosplenial cortex (RSPv). L5 neurons could send axonal branches to targets both in and out of the DMN, broadcasting information to multiple types of downstream targets. Alternatively, specific subclasses of projection neurons may exist that preferentially target either in- or out-DMN targets. To test these possibilities, we used a dual retrograde/anterograde intersectional viral tracing method^53^ to label all the axon branches from neurons defined by whether they project to an in-our out-DMN target. We identified diverse patterns of target specificity in prefrontal, medial, and visual sources, and report two projection neuron subclasses in RSPv that are strongly correlated with the DMN.

## Results

### Voxel and regional spatial maps of the mouse DMN

We started with a dataset of n=40 blood-oxygen-level dependent (BOLD) rsfMRI experiments performed on lightly anesthetized mice^51^ (available online at https://doi.org/10.17632/7y6xr753g4.1, https://doi.org/10.17632/r2w865c959.1). We identified five components using low-dimensional group independent component analysis (ICA). Four matched previously described networks^5^: (1) default mode, (2) somatomotor, (3) lateral cortical, and (4) hippocampal. The fifth component contained cerebellar signal and apparent noise and was not used for further analysis.

To precisely identify the spatial coordinates and brain regions that comprise the DMN and other functional networks, we aligned them to the Allen CCFv3 (**Figure 1a**). The fMRI timeseries data was initially registered to an in-house mouse brain template (~100×100×500μm), which we projected into the CCFv3 reference space using a combination of linear (affine) and non-linear registrations. The parameters from the template transform were used to map the ICA components into the CCFv3 space (100 μm^3^ voxels). Then, we thresholded each ICA component at a z-score = 1. We also include a z-score = 1.7 threshold to help identify “core” DMN structures. We symmetrized each fMRI mask by shrinking to the minimum voxel-wise overlap between the two hemispheres in CCF space, producing 3D spatial masks of four mouse functional networks registered to the Allen CCFv3 at 100 μm resolution (**Figure 1b,c, Figure S1a-c**). A similar network to the DMN ICA component could be produced from the BOLD data using a seed region placed in the anterior cingulate cortex, a core constituent of the rodent DMN^8^ (**Figure S1d**).

**Figure 1.**
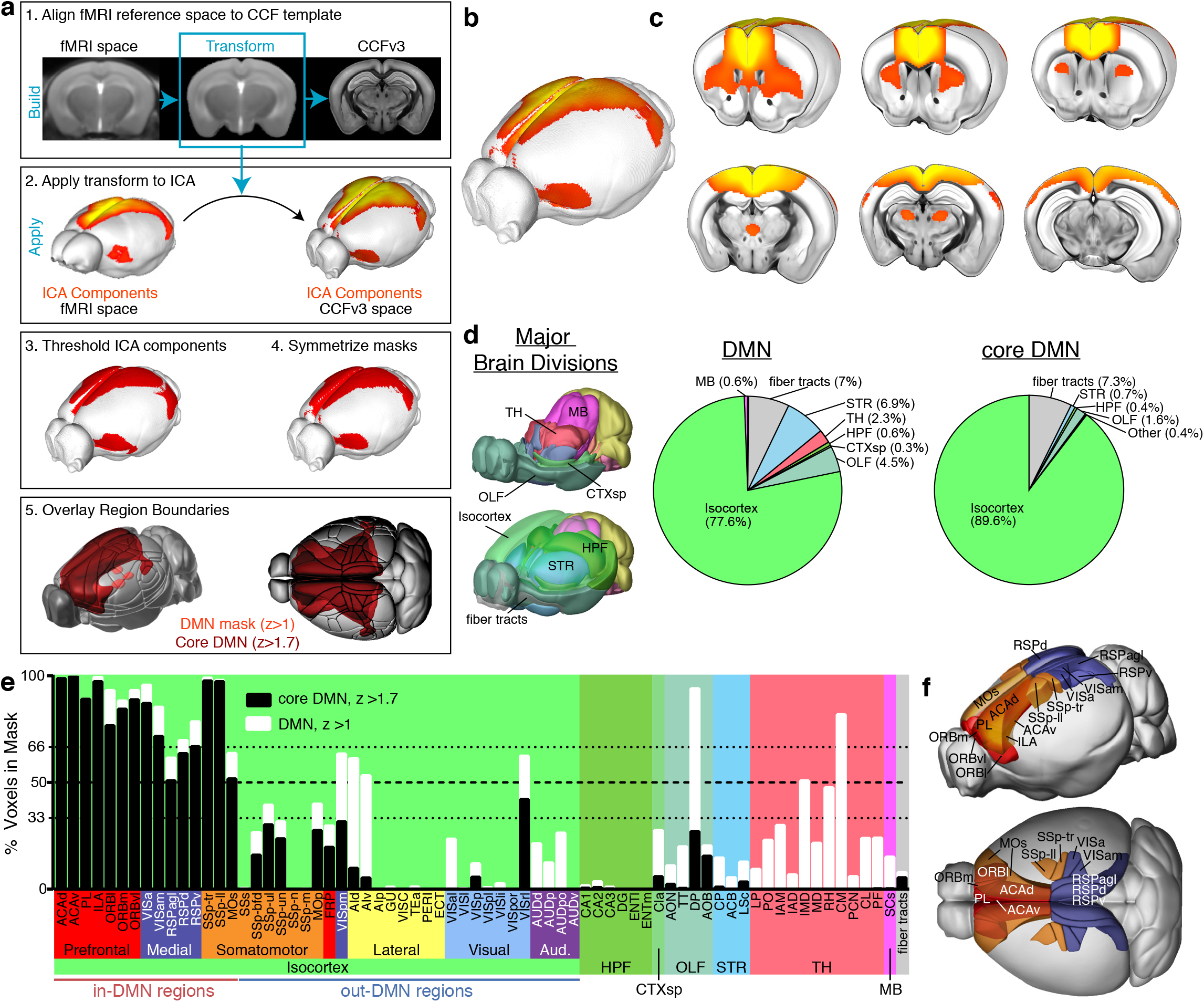
Anatomical identification of DMN structures. **a** Workflow for registering ICA components to the Allen CCFv3: 1. Deformably register in-house fMRI template to CCFv3, 2. Apply the transformation obtained from the templates to the ICA data, 3. Threshold at z=1 for components 1-4 masks and z=1.7 for the DMN core mask, 4. Symmetrize the masks by taking the minimum voxel-wise overlap between hemispheres, 5. Overlay regional boundaries from the Allen Mouse Brain Reference Atlas. **b** 3D projection image of the ICA component corresponding to the DMN overlaid on the CCFv3 template. **c** Serial cutaway images showing coronal sections of the DMN ICA component in CCFv3 space. **d** Percentage of mask voxels in the DMN and core masks that belong to each major brain division. Abbreviations: TH: thalamus, MB: midbrain, OLF: olfactory areas, CTXsp: cortical subplate, HPF: hippocampal formation, STR: striatum. **e** Percent of the voxels that overlap with the DMN mask at both z scores (white bars z>1, black bars z>1.7) shown for all isocortex structures and selected structures in other major brain divisions that overlapped with the DMN and core masks. Thick colored bars under isocortex structures indicate the module affiliation for each cortical region as in Harris et al. 2019. DMN regions were defined as those with >50% of their voxels in both the DMN and core masks. **f** The anatomical location of DMN regions colored by their module affiliation (Prefrontal: red, Medial: blue, Somatomotor: orange).

We used the atlas-aligned resting state network masks to calculate the overlap between all the voxels assigned to the DMN with the structure annotations in CCFv3, starting at the major brain division level. Confirming what was visually obvious and consistent with previous reports, the DMN mask is mostly cortical; 77.6% of the voxels belong to the isocortex. The rest of the DMN mask (22.3%) is comprised of voxels within fiber tracts, striatum (STR), olfactory areas (OLF), medio-dorsal thalamus (TH), hippocampal formation (HPF), midbrain (MB), and cortical subplate (CTXsp, **Figure 1d**). The core DMN mask was even more restricted to the cortex, with 89.6% of its voxels overlapping isocortex, 7.3% in fiber tracts, and less than 3% in other major brain divisions. Overlap of the other three resting state networks with major brain divisions are shown in **Figure S1e-g**. Next, we determined the specific brain regions within these major divisions that contain DMN voxels. We report the percent of voxels in each structure that are part of the DMN and core-DMN masks for 316 regions brain-wide (*i.e.,* the “summary structure set”, **Supplementary Table 1**). Within isocortex, we identified 36 distinct areas with at least one voxel in the DMN mask and 25 areas in the core-DMN mask (**Figure 1e**). Fewer regions (n=1 to 11) were identified within the other major divisions that had any overlap with the DMN masks. Notably, the claustrum (CLA) was the single region in the CTXsp with both DMN and core voxels. In the thalamus, which contained 2% of the total DMN mask, specific structures included several midline nuclei, *i.e.,* central medial (CM), intermediodorsal (IMD), rhomboid (RH) and interanteromedial (IAM). None of these structures were present in the core-DMN mask (white bars only in **Figure 1e**). We chose to conservatively define regions inside the DMN (“in-DMN”) as those with > 50% of their voxels in the core-DMN mask; only sixteen regions passed this threshold (**Figure 1e**, black bars over thick dashed line). These structures were all in the isocortex except one, induseum griseum, a thin, elongated structure that runs along the midline adjacent to anterior cingulate cortex (ACA) and cannot be individually resolved. Of note, four cortical regions had >50% of their voxels inside the z=1 but not the z=1.7 DMN mask: posteromedial and rostrolateral visual areas (VISpm and VISrl) and the dorsal and ventral insular cortex (AId and AIv). These regions were considered “out-DMN regions” for subsequent analyses. The core mask was used only to define DMN structures; all subsequent analyses used the z=1 DMN mask.

We noted a very strong resemblance between the spatial distribution patterns of the DMN masks and cortical network modules previously derived from our structural connectivity analyses^25^. Here, we grouped all isocortex structures by their module membership, showing that most of the in-DMN structures (n=12 of 15) belong to two of the six modules; prefrontal and medial (**Figure 1e**). Exceptions include the trunk and lower limb regions of somatosensory cortex (SSp-tr and SSp-ll), and secondary motor cortex (MOs). Conversely, most regions belonging to these two modules were defined as in-DMN structures, with two exceptions; frontal pole (FRP) and VISpm (borderline for “in-DMN” as mentioned above). Spatial locations for all 15 in-DMN regions are shown in angled lateral and dorsal views of the CCF, colored by module membership (**Figure 1f**). The DMN can thus be defined most specifically by the voxel-based mask, alternatively by a set of 15 anatomically parcellated regions, and approximately by the combination of the prefrontal and medial modules.

### Preferential region-to-region connectivity in the DMN

Segregation of the majority of in-DMN regions into the two network modules suggests that cortico-cortical connections inside the DMN are stronger than connections made to cortical areas outside the DMN. So, we directly quantified the degree to which in-DMN regions preferentially connect with each other by examining projection data from a set of 300 anterograde viral tracer experiments distributed across the cortex (**Figure 2a**). This dataset includes injections into wild type mice^24^ (WT) and Emx1-Cre driver mice, which label all cortical projection neuron classes^25^. We also included experiments in the Rbp4-Cre_KL100 line (Rbp4), which preferentially labels layer 5 (L5) output neurons, based on our previous work showing a significant correlation in connection strengths and no difference in targets reached with WT experiments^25^.

**Figure 2.**
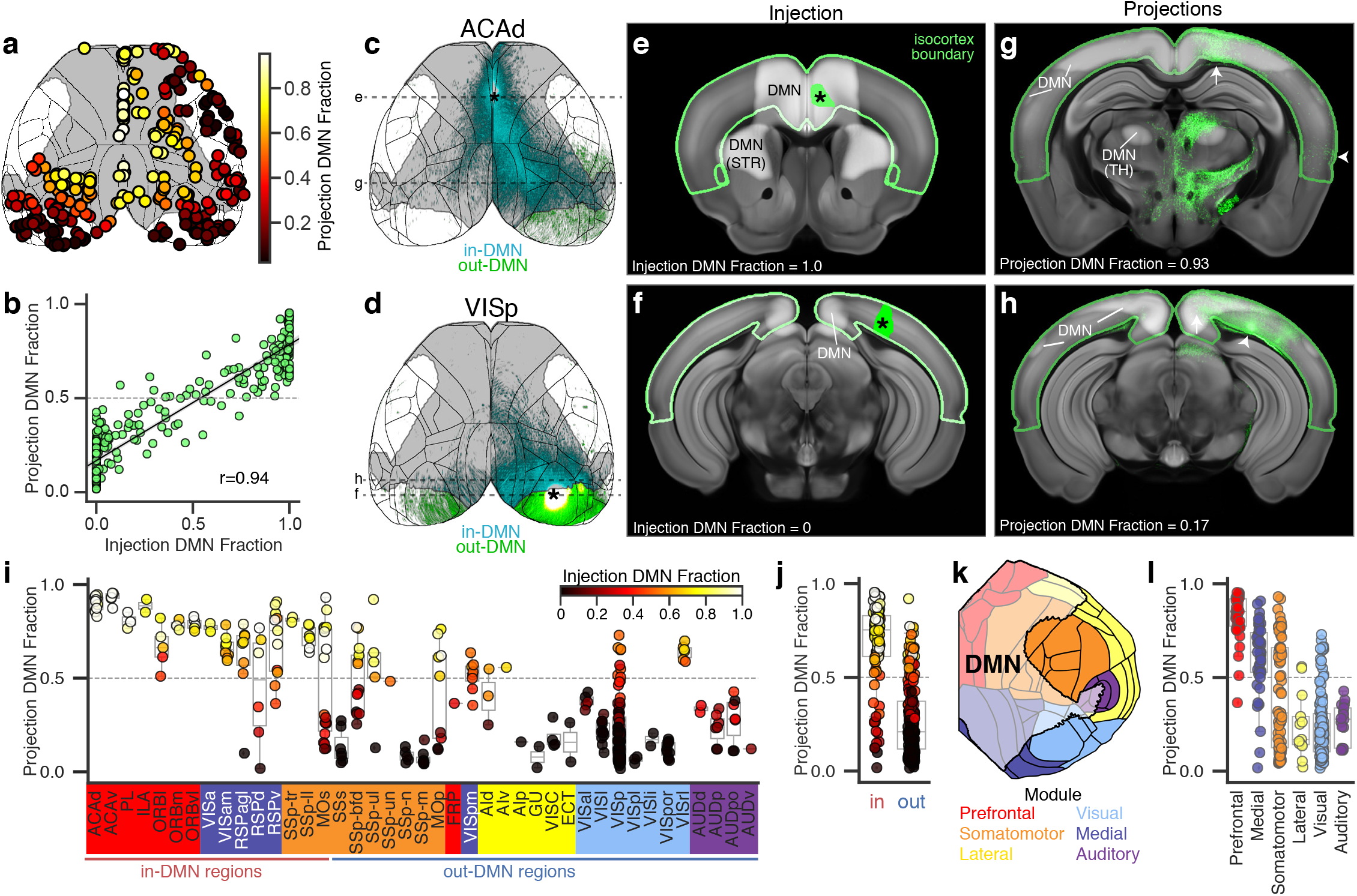
DMN regions preferentially project to other DMN regions. **a** Top-down cortical projection map showing the location of the 300 injection experiments in WT, Emx1-Cre, and Rbp4-Cre mice used to quantify preferential DMN connectivity. The boundary of the DMN mask is shown in gray and cortical structure boundaries in black. Colormap indicates the percent of cortical projections inside the DMN mask for each experiment. **b** Scatterplot and linear fit for the percent of cortical projections inside the DMN as a function of the percent of the injection inside the DMN for each of the 300 cortical injections shown in a. r = Pearson correlation. **c,d** Cortical projection images showing an injection experiment that was completely inside the DMN mask (ACAd, **c**), and an injection located completely outside the DMN mask (VISp, **d**). Asterisks indicate the approximate centroid of the injections. In-DMN projections are pseudocolored cyan and out-DMN projections green. Full image series available online at http://connectivity.brain-map.org/projection/experiment/cortical_map/[insert experiment ID]. Experiment ids: ACAd 112458114, VISp 100141219. Dashed lines in **c** and **d** show the rostral-caudal position of coronal template sections in **e-h**. **e,f** Single coronal image sections through the center of the ACAd (**e**) and VISp (**f**) injection sites showing segmented injection pixels (green) overlaid on the corresponding coronal section from the CCFv3 template. **g,h** Single coronal sections at the approximate rostral-caudal position with the highest projection density showing segmented projection pixels (green) overlaid on the corresponding CCFv3 template section. Arrows indicate in-DMN projections and arrowheads indicate cortical projections outside the DMN. **e-h** White overlay: DMN mask, green edges: isocortex boundary. Portions of the DMN mask overlapping striatum (STR) and thalamus (TH) are labeled in **e** and **h**. **i** Fraction of cortical projections inside the DMN for each of the 300 experiments shown in a grouped by their injection source. Sources are arranged by module and DMN membership. Boxplots show the range of values for all experiments in each source, and individual points for each experiment are colored by the percent of their injection that is inside the DMN mask. Note that some sources have injections both inside and outside the DMN. **j** The same points as in i with sources grouped by in-DMN and out-DMN regions. **k** Cortical flat map with regions colored by their module affiliation. White overlay shows the DMN mask borders. **l** The same points as in f grouped by module affiliation.

First, we performed voxel-level analyses. For each experiment, we compared the fraction of the injection site inside the DMN mask with the fraction of segmented cortical projections within the DMN mask. We observed a significant positive correlation between these measures (r = 0.94, p<0.001, **Figure 2b**). Axonal projections labeled from two example injections, one 100% inside and one 100% outside of the DMN, are shown in top down cortical surface views (**Figure 2c,d**) and as overlays on virtual coronal slices from the CCF average template with the DMN mask (**Figure 2e-h**) to illustrate their overlap with the DMN voxel mask (gray or light gray color in **Figure 2c-h**).

The DMN mask is spatially continuous as it stretches from anterior to posterior along the midline and across parietal to posterolateral cortical regions (**Figure 1b,c**). So, it is not altogether surprising to find that injections in DMN regions send a higher fraction of their projections to other DMN regions, given that connection density is well-known to decrease as a function of distance^24,54,55^. To quantify the extent to which preferential intra-DMN connectivity can be accounted for by spatial proximity, we applied a general linear model (GLM) to derive a distance coefficient and a DMN coefficient for each experiment. The model fit for two experiments is shown in **Figure S2a,b**, and the centroid location for all 300 experiments is shown in **Figure S2c**, colored by DMN coefficient. Distance coefficients were negative or close to zero and had a low inverse correlation with the fraction of the injection inside the DMN mask (r=−0.16, p=0.005 **Figure S2d**). The DMN coefficient, however, was positively and significantly correlated with the fraction of the injection in-DMN (r = 0.60, p<0.001, **Figure S2e**). These results show that even after accounting for distance effects on connectivity strengths, DMN regions still send more projections to other DMN regions. In some cases, the DMN coefficient can provide insight into which regions should be considered DMN structures. Dorsal agranular insula (AId), for example, had a slightly negative DMN coefficient, supporting its exclusion from the list of DMN structures, but both the rostrolateral visual area (VISrl, a subdivision of the posterior parietal association area, PTLp) and ventral agranular insula (AIv) had slightly positive DMN coefficients (**Figure S2c**).

We wondered whether the tendency for regions inside the DMN to send more projections inside that network is a unique property of the DMN or a more general property of resting state networks. To test this, we calculated “mask” coefficients and distance coefficients for the other ICA components in our fMRI dataset. In all cases, there was a linear relationship between the fraction of an isocortex injection that overlapped the functional mask and the fraction of its cortical projections inside that functional mask (**Figure S2f,g**). For the hippocampal network, this relationship also held for hippocampal injection experiments (**Figure S2g**). In fact, the correlations were stronger for the other three networks than for the DMN suggesting that the DMN may be *less* preferentially connected than other functional networks, a finding consistent with the polymodal structure of this distributed set of regions^56^.

Next, we performed region-level analyses by grouping the 300 isocortex injections into in- or out-DMN brain regions, as defined in **Figure 1e**. Experiments with injections that spanned multiple source regions were categorized as in- or out-DMN based their primary source region, i.e. the region that contained the majority of the injection volume^57^. As seen at the voxel-level, in general, injections into in-DMN regions had a higher fraction of DMN projections than injections into out-DMN regions (**Figure 2i,j**). We again used a GLM to separate the effects of distance and DMN connectivity, finding that experiments in most in-DMN structures indeed had positive DMN coefficients, although a few were zero or negative (**Figure S2h**). For example, the lateral orbital area (ORBl) sent fewer than expected projections to the DMN once distance was taken into account in one experiment (indicated by its negative DMN coefficient).

Across regions, the fraction of a given injection volume inside or outside the voxel-level DMN mask was more predictive of the fraction of projections inside the DMN than whether or not the source region was assigned to a DMN structure based on anatomical parcellations (see colormap for individual points in **Figure 2i** and compare **Figure 2b** with **Figure 2j**). In fact, in several locations the boundary of the DMN mask does not align with anatomical boundaries (**Figure 2k**). For example, ~ 50% of the voxels belonging to secondary motor cortex (MOs) are in the core-DMN and ~60% in the DMN mask (**Figure 1e**), and injections into the DMN portion of MOs had more DMN projections while injections outside the DMN in MOs had fewer DMN projections (*n.b.* the bimodal distribution in **Figure 2i** and see **Figure S3**). We also summarized the fraction of projections in the DMN mask by cortical module (**Figure 2l**). Most injections in prefrontal and medial modules had >50% of their cortical projections inside the DMN mask. Conversely, nearly all injection experiments in lateral and auditory modules had <50% of their cortical projections inside the DMN. These results were still consistent when accounting for distance (**Figure S2i-j**). A summary of the cortical regions and projections that intersect the core-DMN mask is provided in **Supplementary Table 2**. These results confirm that DMN regions are more strongly connected to each other than they are to regions outside the DMN and suggest that the DMN may cover functionally segregable sub-portions of previously defined anatomical structures.

### Preferential connectivity between DMN regions by layer and projection cell class

The polymodal nature of the DMN^56^ and its central role as integrator of between-network communication^13,29,31^ mean that although cortical DMN regions may send most of their projections to other cortical DMN regions, they also target regions outside the DMN. We hypothesized that distinct cell classes within these regions might contribute to this target specificity, with one class carrying in- and one class carrying out-DMN projections. Cortical regions are comprised of distinct excitatory neuronal types organized by layers and long-distance projection patterns: intratelencephalic (IT) in L2-L6, pyramidal tract (PT) in L5, and corticothalamic (CT) in L6^38,58^. Cortico-cortical projections originate mostly from IT neurons in L2/3 and L5^24^. We reported recently that L5 neurons have the most extensive cortical projections; cells in other layers within the same region project to a subset of the L5 cortical targets^25^. Thus, one possibility is that connections to both in- and out-DMN areas are made by L5 cells, and that the stronger in-DMN connection weights comes from additional cells in other cortical layers projecting specifically to DMN targets.

To assess for layer-specific projection patterns related to in-DMN regions, we started with a set of spatially-matched viral tracing experiments in 15 Cre and WT lines, “anchored” by the L5-selective Rbp4-Cre_KL100 line^25^. We further curated this dataset so that each Rbp4 anchor group contained in-DMN or out-DMN injection experiments, but not both (see methods). This resulted in a set of 350 experiments across 42 spatially matched groups. Each group contained at least one experiment from a Cre line targeting L2/3, L4, L5 IT, L5 PT, and L6 CT, plus a wild type (WT) or Emx1-IRES-Cre (Emx1) injection. We focused primarily on experiments in Cre lines selective for L2/3 and L5 IT cells due to their more extensive intracortical projections (**Figure 3**); but data and analyses from the L4 IT, L5 PT, and L6 CT experiments are also provided (**Figure S4**). Experiments were distributed across the cortex and the DMN mask (n=11 of 15 in-DMN sources, 13 of 28 out-DMN sources, **Figure 3a,d**). Cortical projections from the group of experiments in anteromedial visual cortex (VISam, a DMN region) are shown to illustrate the different layer-specific cortical projection patterns (**Figure 3b**). As we previously reported, WT and Rbp4 experiments have the most extensive cortical projection pattern. Notably, L2/3 projections visibly overlap those from L5, but target less of the cortex, particularly in the contralateral hemisphere^25^ (**Figure 3b**). We plotted the fraction of the Cre line injection in the DMN against the fraction of its cortical projections in the DMN (**Figure 3c**). Like the WT experiments (**Figure 2b**), these values are also significantly correlated for all Cre lines (r=0.91-0.96, all p<0.001). L2/3 experiments inside the DMN (injection DMN fraction > 0.5), had a significantly higher fraction of their projections in the DMN than L5 experiments inside the DMN (p=0.007, **Figure 3c,** inset). Notably, cortical projections from L4 IT, L5 PT, and L6 CT neurons are much more restricted and predominantly target nearby regions^25^ (see also **Figure S4b**). Experiments from these layers and classes had less negative distance coefficients (**Figure S4d**) compared to WT, L2/3 and L5 IT experiments, reflecting the limited spatial spread of their projections. The fraction of DMN projections was significantly higher for L2/3 compared to L5 experiments, for in-DMN, but not out-DMN, sources (p=0.003, **Figure 3d,e**). In fact, the median fraction of in-DMN projections for L2/3 in-DMN sources was nearly 100% (0.91); in other words, almost all of the projections from L2/3 in-DMN neurons are to other DMN regions. The median fraction for L5 output was 0.73, meaning that ~ 30% of the L5 projections are outside of the DMN mask.

**Figure 3.**
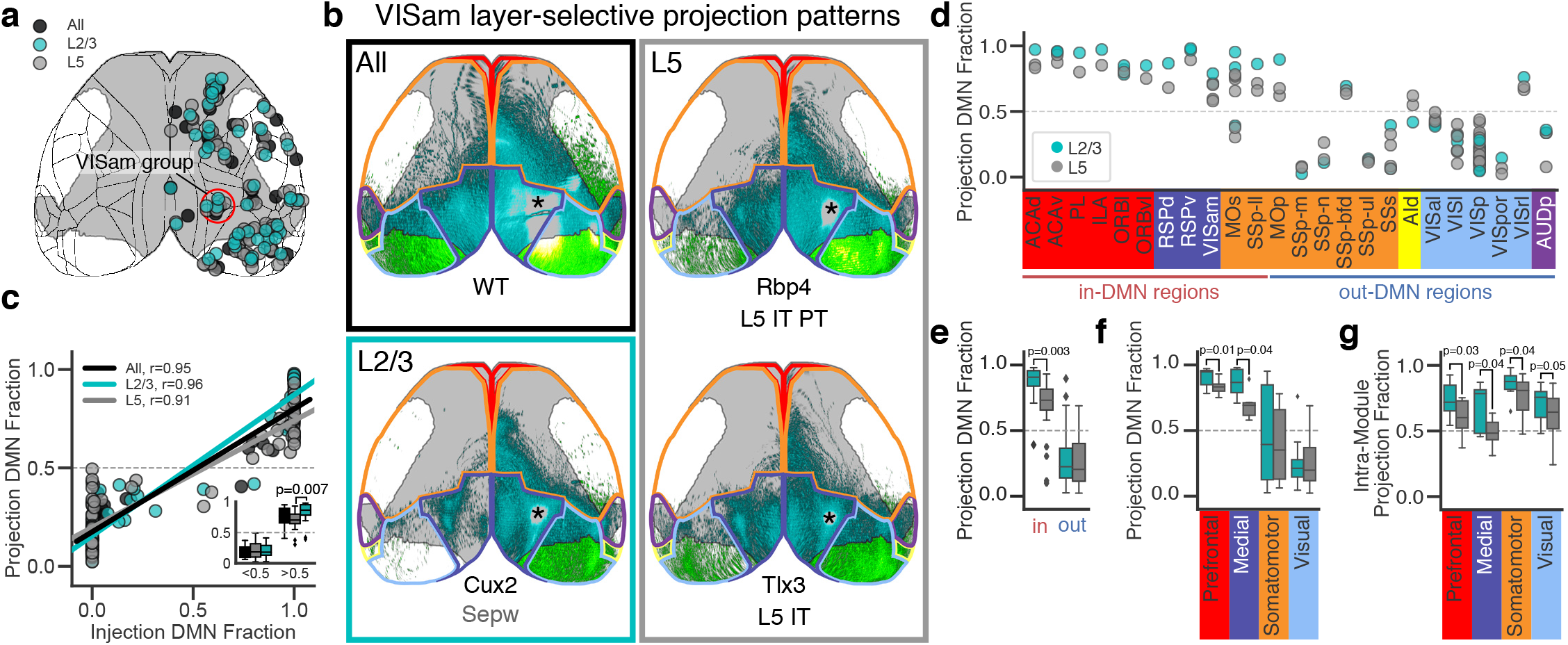
Layer 2/3 neurons have more intra-DMN and intra-module projections than layer 5 neurons. **a** Top-down cortical map showing the location and relative size of matched injection experiments in WT and Emx1-IRES-Cre mice (black, n=41), L2/3 Cre-driver lines (cyan, n=46), and L5 IT Cre driver lines (gray, n=76). Red circle indicates the injection centroids belonging to the anteromedial visual cortex (VISam) group, shown in **b** and **Figure S4**. Gray overlay shows the DMN mask borders. **b** Top down cortical projection of matched experiments in a WT mouse, a Cux2-Cre mouse selective for L2/3 neurons, and Rbp4-Cre and Tlx3-Cre mouse lines selective for L5 IT neurons. Experiments in Sepw1-Cre mice were also used for L2/3-selective projections, but there was no Sepw1 injection in VISam. In-DMN projections are pseudocolored cyan and out-DMN projections are green. Asterisk indicates the approximate location of each injection centroid. Gray overlay shows DMN mask and colored lines indicate module boundaries. Full image series available online at http://connectivity.brain-map.org/projection/experiment/cortical_map/[insert experiment ID]. Experiment IDs: WT 100141599, Cux2 184167484, Rbp4 159753308, Tlx3 297233422. **c** Scatterplot and linear fit for the percent of cortical projections inside the DMN as a function of the percent of the injection inside the DMN for each of the 163 cortical injections shown in **a**. Inset shows the fraction of DMN projections for injections binned by the fraction of the injection polygon inside the DMN mask. **d** Fraction of cortical projections inside the DMN for the L2/3 and L5 experiments shown in a grouped by their Rbp4 anchor group source (see methods and Harris et al. 2019) and colored by layer. Sources are arranged by module and DMN membership. **e,f** Boxplots showing the distribution of the points grouped by in-DMN and out-DMN sources (**e**), and module (**f**). L2/3 injections inside the DMN had a higher fraction of their projections inside the DMN than spatially matched L5 injections for both prefrontal and medial sources. **g** Boxplots showing the fraction of intra-module projections for experiments in L2/3 and L5 selective Cre lines. Only modules with more than one injected source are shown. **c,e,f,g** Boxes=IQR, whiskers=1.5×IQR, p-values were determined by multi-way ANOVA followed by Tukey’s post hoc test with correction for multiple comparisons.

Previously, we showed that projections arising from L2/3 IT neurons differ most from L5 IT neurons in the contralateral hemisphere^25^. Specifically, L2/3 IT neurons have more restricted contralateral projections, primarily to homotypic regions. Given that the DMN mask is symmetric across both hemispheres, we speculated that the higher fraction of projections in the DMN from L2/3 vs L5 might be simply explained by this difference in contralateral projections. However, when we removed the contralateral projections and repeated these analyses, the relative values between groups did not change (**Figure 3c** and **S4c-e** vs **S4f-h**).

When grouped by module, injections in the prefrontal and medial modules both had significantly more DMN projections from L2/3 compared to L5 (**Figure 3f,** p=0.01 for prefrontal module, p=0.04 for medial module). Since the DMN is mainly composed of these two modules, this difference may reflect an overall tendency for L2/3 cells to have mostly intra-module projections compared to L5, independent of the module of interest. We assessed for this by measuring the fraction of intra-module projections for each of the four modules with n>1 source in our dataset (prefrontal, medial, somatomotor, visual) and found that in all cases L2/3 neurons had a higher fraction of intra-module projections than L5 (**Figure 3g**). This is also the case for the single source in the auditory module (AUDp), but not for the single lateral source in this dataset (AId, **Figure 3d**). Together, these results show that DMN target regions receive input from both L2/3 and L5 cells in DMN source regions, but that projections outside the DMN mask from DMN sources arise predominantly from L5 while L2/3 cells primarily project within the DMN.

### Preferential connectivity between DMN regions by target-defined neuron class

Cre lines are useful for labeling cell classes, but most contain a heterogeneous mix of cell types as defined based on transcriptomic, morphological, and physiological properties^41–43,59^. We showed using Cre line tracing data that, at the population level, L5 cells in DMN regions project both in and out of the DMN. Where a cell projects is also a defining feature of a “type”. Recent retrograde and single cell projection data supports the existence of IT and PT neuron subclasses that project to a subset of a region’s total target set in non-random combinations^60–62^. So, we hypothesized that L5 projection neuron types may co-exist in DMN regions that preferentially target either in- or out-DMN regions. To test this, we used an intersectional viral tracing approach to label all the axons of target-defined (TD) projection cell types^53^. We injected a retrogradely-transported canine adenovirus encoding Cre (CAV2-Cre^63,64^) into a “primary target” region and a Cre-dependent adeno-associated virus (AAV-FLEX-EGFP) into a “source” region in Ai75 reporter mice^49^ that express nuclear-localized tdTomato (nls-tdT) in the presence of Cre **Figure 4a,b**. Thus, local and long-distance neurons with axon terminals in the primary target site will express Cre following retrograde transport and are visible brain-wide as nls-tdT+. In the source region injected with AAV, co-infected cells that express Cre (and nls-tdT) will also express EGFP. EGFP labels the entire cell (cell body and axons), allowing visualization of projections to both the primary target (CAV2-Cre injection site) as well as all other targets (secondary targets, **Figure 4b**). We used the same serial two-photon tomography (STPT) imaging and informatics pipeline developed for the Allen Mouse Connectivity Atlas (MCA^24,25,57^) to quantify brain-wide EGFP fluorescence in the intersectional viral tracing experiments.

**Figure 4.**
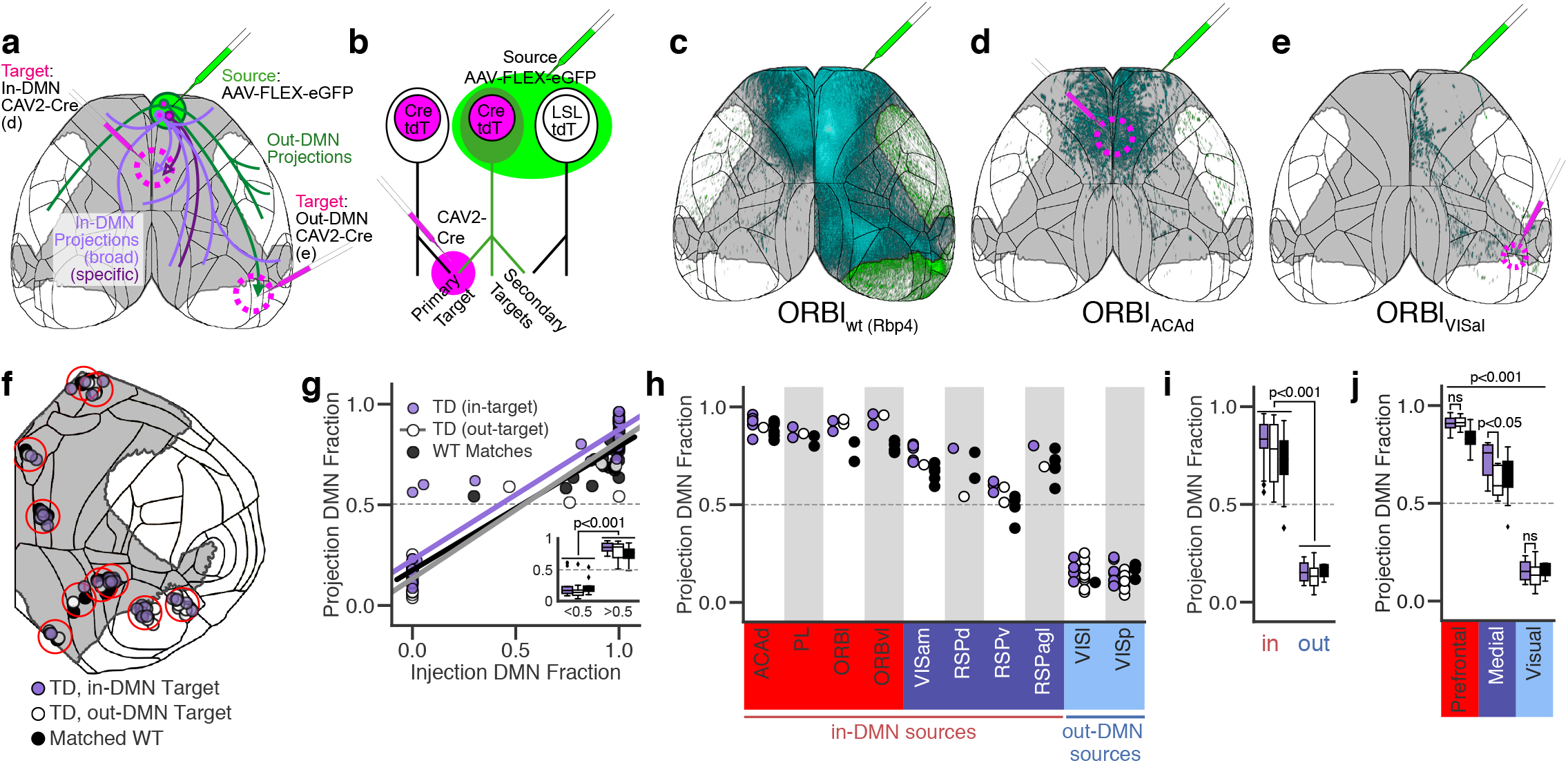
Target-defined projection mapping differentiates in-DMN and out-DMN projections for sources in the medial module. **a** Experimental design for target-defined (TD) projection mapping. For each experiment, a “source” was injected with Cre-dependent adeno-associated virus (AAV-FLEX-EGFP) and a retrograde transported virus encoding Cre (CAV2-Cre) was injected into a “target”. Cells in the source that have axon terminals in the target are infected with both viruses allowing visualization of their brain-wide projections. We hypothesize that in-DMN sources paired with in-DMN targets will have more in-DMN projections (purple cells) than in-DMN sources paired with out-DMN targets (green cell). Cells labeled in TD experiments may project to only a few (specific) or many (broad) of the targets observed in WT experiments. **b** Method and terminology for TD experiments. Each experiment has one source injection and one target injection. Cells that project to the primary target (CAV2-Cre injection site) can be visualized brain-wide by their nuclear tdTomato (tdT) expression, produced by the viral-expressed Cre from the floxed nuclear tdT gene in the Ai75 mouse line (LSL tdT). Cells that are also infected with the AAV-FLEX-EGFP source virus express cytosolic eGFP, allowing visualization of their brain-wide projections to other targets (secondary targets). Cells infected with the AAV-FLEX-EGFP source virus but not the CAV2-Cre target virus are not fluorescent (right) **c** A mesoscale tracing experiment in an Rbp4-Cre mouse from the Allen Mouse Connectivity Atlas with source cells in lateral orbital cortex (ORBl) has extensive cortical projections. **d,e** Top-down cortical projection images from TD experiments with a source injection in ORBl paired with an in-DMN target, dorsal anterior cingulate area (ACAd, **d**) or an out-DMN target, anterolateral visual area (VISal, **e**). The DMN mask is shown in gray, and in-DMN projections are pseudocolored cyan. **d** and **e** were both left hemisphere injections, but they are flipped to the right hemisphere to facilitate comparison with the Rbp4 experiment in **c**. **c,d,e** Full image series available online at http://connectivity.brain-map.org/projection/experiment/cortical_map/[insert experiment ID]. Experiment IDs: ORBl_Rbp4_ 156741826, ORBl_ACAd_ 571816813, ORBl_VISal_ 601804603. **f** Cortical flat map showing the location of the ten groups of experiments with matched injection sites (red circles) and at least one experiment of each type (TD in-target, TD out-target, WT). The boundary of the DMN mask is shown in gray and cortical structure boundaries in black. **g** Scatterplot and linear fit for the fraction of projections inside the DMN as a function of the fraction of the injection in the DMN for each of the three types of experiments. Inset shows the fraction of DMN projections for injections binned by the fraction of the injection polygon inside the DMN mask. **h** Boxplots showing the fraction of projections inside the DMN for the groups of injection-matched experiments shown in f grouped by their source region. **i** Boxplots showing the distribution of DMN projection fractions for each experiment type for in-DMN and out-DMN sources. **j** Boxplots showing the distribution of DMN projection fractions for each of the three modules that contained a group of injection matched experiments. **g,i,j** boxes=IQR, whiskers=1.5×IQR, p-values were determined by multi-way ANOVA followed by Tukey’s post hoc test with correction for multiple comparisons.

TD experiments are named by their primary source with a subscript indicating their primary target, i.e. an ACAd_RSPv_ experiment has an AAV-FLEX-EGFP source injection into anterior cingulate area, dorsal part (ACAd) and a CAV2-Cre target injection into ventral retrosplenial cortex (RSPv), so all the cells expressing EGFP in that experiment have cell bodies in ACAd and project to RSPv. This naming convention only incorporates the primary injection structure for both the source and target, but injections often covered two or more regions. When there was a nearly equal split between two target regions, we indicated both targets in the subscript (e.g. ACAd_RSPv/RSPd_). The full source/target coverage for each experiment is available in the online dataset (http://connectivity.brain-map.org; filter for Tracer Type: Target EGFP). TD cells labeled by intersectional tracers may project to many (broad) or few (specific) targets (**Figure 4a**). Importantly, CAV2-Cre has a known tropism bias towards L5 over L2/3 neurons^65^. Although this bias is expected to be in our favor in these experiments (given the L5 IT focus), we want to emphasize that we are likely missing data from L2/3 IT cell types.

The target location for any given source was chosen based on analyses of projections labeled in WT, Emx1, or Rbp4 mice. For example, L5 IT axons originating from ORBl project widely to multiple cortical targets (**Figure 4c**), including ACAd (in-DMN) and the anterolateral visual area (VISal, out-DMN). To determine what secondary targets (if any) are reached by ORBl neurons that target ACAd (ORBl_ACAd_), we injected AAV-FLEX-EGFP into ORBl and CAV2-Cre into ACAd (**Figure 4d**). Visually, ORBl_ACAd_ cortical projections appeared to be a subset of the total ORBl L5 projection pattern (compare to **Figure 4c**); axons were densest in the rostral DMN in both hemispheres and also targeted lateral cortical areas outside the DMN. In contrast, an injection at the same location in ORBl paired with a different primary target, VISal (ORBl_VISal_), revealed a different subset of the ORBl L5 projections (**Figure 4e**). High-resolution images from these source and targets are shown in **Figure S5**. These results show that TD projection mapping can reveal types of intracortical projection neurons with distinct multi-area targeting patterns. We performed 183 TD projection mapping experiments in total for 35 sources, 15 in the isocortex and 10 in the thalamus and hippocampal formation. Replicate injections (n=2-9) were included for many source-target pairs, but some had only n=1 (see methods). Data (including high resolution images, segmentation masks, and informatically-derived projection weights) are available through the Allen Mouse Connectivity Atlas data portal and associated API and SDK (http://connectivity.brain-map.org; filter for Tracer Type: Target EGFP).

We wanted to identify TD experiments with *limited or specific*, as opposed to *broad,* projection patterns (see **Figure 4a**), based on the assumption that limited TD projection patterns are more likely to be formed by a distinct subclass of target-defined projection neurons. TD experiments had a smaller volume of infected cells in the source region than WT or Cre experiments (**Figure S6a**), and small injections are expected to contact fewer targets than larger injections, particularly if the larger injection includes infection across borders into secondary sources. We wanted to be certain that each TD experiment could reasonably be expected to represent a subset of the WT projections in any experiment to which it was compared, so we first developed a method for systematically identifying matched experiments. In brief, four criteria were applied: (1) primary injection structure was the same, (2) if the primary injection structure was < 60% of the injection volume in either experiment, the secondary injection structure (i.e. the structure containing the second-highest fraction of the injection volume) was also the same, (3) injection centroids were ≤ 800 μm apart, and (4) the Dice Coefficient (to measure spatial overlap) was > 0.05.

Using these criteria, we identified ten sets of injection-matched cortical experiments comprised of at least one experiment per type: (1) TD with an in-DMN target, (2) TD with an out-DMN target, and (3) a WT, Emx1, or Rbp4 L5 experiment from the MCA (see methods). To classify a TD experiment as either an “in-DMN” or “out-DMN” target, we calculated the percent of the CAV2-Cre injection polygon that overlapped with the DMN mask and used a threshold of 50% (>50% = “in-DMN targets”, <50% = “out-DMN”). This curated set includes data for ten cortical areas with 105 total experiments (63 TD and 42 WT); 8 (of 15) in-DMN and 2 out-DMN sources. Experiment IDs are provided in **Supplementary Table 3**. Injection locations are plotted on a cortical flat map in **Figure 4f.** For TD experiments, we report the fraction of projections inside the DMN mask after computationally removing GFP segmentation in the CAV2-Cre injection polygon (see methods and **Figure S7**).

First, we performed the same voxel-level correlation analysis done for WT and Cre line data (**Figure 2b** and **Figure 3c**), again finding significant correlations between the fraction of a source injection and the fraction of cortical projections in the DMN for both in- and out-DMN TD experiments (r=0.94 TD in-target, 0.96 out-target, 0.95 matched WT, all p<0.001, **Figure 4g**). Next, we compared the fraction of projections in the DMN by source. Overall, there was no difference in the fraction of in-DMN projections for sources paired with in-DMN targets compared to those paired with out-DMN targets (**Figure 4h, i**), but there was a considerable amount of variability in the DMN projection fraction for in-DMN source experiments, particularly when paired with out-DMN targets (white boxplot for in-DMN sources in **Figure 4i**). Grouping the sources by module highlighted an important difference between prefrontal and medial sources (**Figure 4j**). Sources in the prefrontal module sent a higher fraction of their projections to the DMN than sources in the medial and visual modules regardless of whether they were paired with an in-DMN or out-DMN source. Sources in the medial module, however, had a significantly higher fraction of projections inside the DMN when paired with in-DMN compared to out-DMN targets (p<0.05). The high fraction of DMN projections from prefrontal sources regardless of their primary target is consistent with a hub-like configuration of this polymodal region, while the higher in-DMN fraction for medial sources paired with in-DMN targets compared with out-DMN targets implies the presence of separate in-DMN and out-DMN projecting cell classes in medial regions.

### Target-defined regional projection patterns vary by source and module

The previous analyses were based on measuring projections across the entire DMN mask. However, we noted that unique subsets of targets were labeled in some TD experiments compared to their matched WT projections (**Figure 4**). Consistent with this, TD experiments project on average to significantly fewer targets compared to their matched WT experiments (n=33±17 targets per TD experiment vs. 63±21 per WT experiment above threshold, p<0.001, Student’s t-test **Figure 5a**)

**Figure 5.**
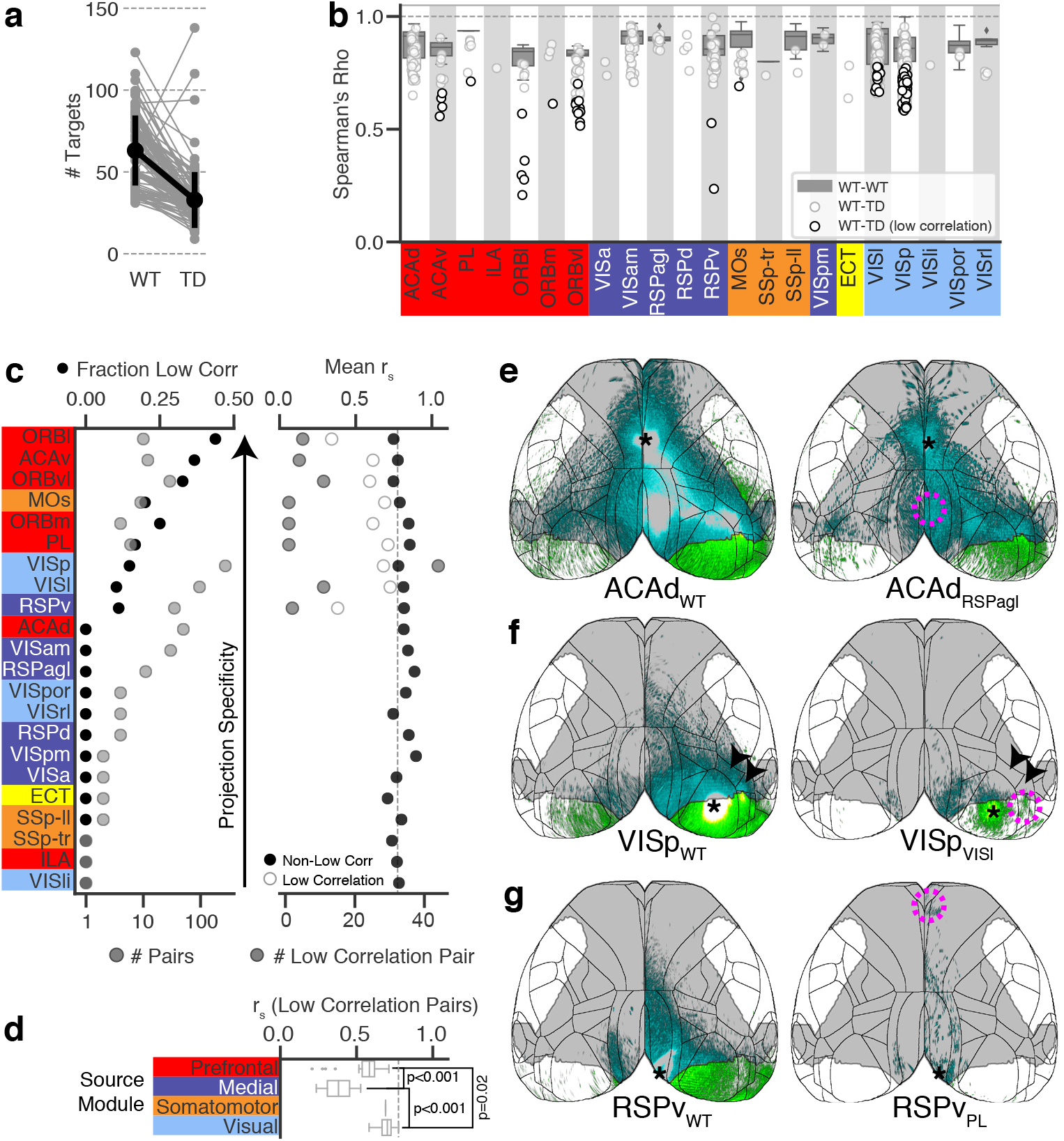
Quantitative comparisons between Target-Defined and WT injection matched pairs. **a** Number of ipsilateral and contralateral targets with normalized projection volume (NPV) > threshold (10^−1.5^) inTD injection experiments compared with the average number of targets above threshold in their WT matches. With only five exceptions, TD experiments contacted fewer targets than their WT matches, both ipsilaterally and contralaterally. **b** Distribution of Spearman’s Rho (r_s_) for WT – WT injection matched pairs (gray boxplots, box = IQR, whiskers = 1.5×IQR) and WT – TD injection matched pairs (white points). Points with black borders are “low correlation” pairs (r_s_ below the 95% prediction interval). Note that the model did not depend on source identity, so six sources had at least one WT – TD match but no WT – WT matches: ILA, ORBm, VISa, RSPd, ECT, and VISli. **c** Sources with TD injections ranked by projection specificity. (left) top axis (black points): Fraction of pairs in each source that were considered low correlation. bottom axis (gray points): Number of experimental pairs for each source. (right) top axis (black and white points) Mean r_s_ value for low correlation WT – TD pairs (white) and non-low correlation WT – TD pairs (black) in each source. bottom axis (gray points): Number of low correlation WT – TD pairs in each source. **d** Distribution of r_s_ values in low correlation pairs grouped by module. WT – TD pairs in the prefrontal module had lower r_s_ values than pairs in the medial and visual module, and pairs in the medial module had lower r_s_ values than pairs in the visual and prefrontal modules. Box = IQR, whiskers = 1.5×IQR. **e-g** Top down cortical projections of matched WT (left) and TD (right) experiments with injections in ACAd (**e**), VISp (**f**), and RSPv (**g**). Asterisks indicate the approximate centroid of each AAV source injection and dashed magenta circles indicate the approximate area of the CAV2-Cre target injection. Full image series available online at http://connectivity.brain-map.org/projection/experiment/cortical_map/[insert experiment ID]. Experiment IDs: ACAd_WT_ 146593590, ACAd_RSPagl_ 607059419, VISp_WT_ 100141219, VISp_VISl_ 515920693, RSPv_WT_ 112595376, RSPv_PL_ 623838656. Arrowheads in **f** show projections in VISrl and VISal that are present in the WT experiment but absent from the TD experiment.

Next, we calculated the Spearman correlation coefficients (r_s_) on measured connection strengths between all pairs of injection-matched WT experiments and for TD experiments with their WT matches. The set of WT-TD pairs had lower r_s_ values than WT-WT pairs (WT-WT: 0.86±0.07, WT-TD: 0.78±0.10, p<0.001, student’s t test). We wanted to use the r_s_ value as a metric for how much a TD experiment’s projection patterns differ from the WT pattern (i.e. how limited the TD experiment’s projections are), but even though we used our match criteria to identify injection-matched pairs, the size difference between WT and TD injection volumes remained an experimental confound (**Figure S6b**). We therefore constructed a model to predict the expected Spearman correlations between two WT experiments based on the volume of the smaller injection in the pair, the distance between the two centroids, and the overlap between the two injection sites (**Figure S6b-d**, see methods). The model was based on a large set of WT replicates (n=1-429 pairs per source, median=5, black points, **Figure S6e**). As expected, the model predicts lower correlation values for smaller injection volumes (**Figure S6e**, black line). We also used the model to calculate the 95% prediction interval for r_s_ given the injection volumes, centroid distances and overlap ratio of each pair of WT-TD experiments (**Figure S6e**, gray line). We used the lower value of this interval as a lower bound to decide whether a given pair of injections was *less correlated* than expected for replicate injections. We called these “low correlation” pairs.

We examined the distribution of Spearman correlation coefficients for all WT-TD pairs by source structure (**Figure 5b**, low correlation pairs have dark borders). Observed correlation values were outside the 95% prediction interval in 21% of all WT-TD pairs (43 pairs above, 80 pairs below). The number of low correlation pairs varied by source region; e.g. n=7 in the ventral part of ACA (ACAv), but n=0 in ACAd, despite ACAd having more experiments (n=50 vs. 12 pairs). Indeed, every source in the prefrontal module had at least one low correlation pair, except for ACAd and the infralimbic area (ILA), which had only one pair. In the medial, somatomotor, and visual modules, low correlation pairs were observed in only one or two sources out of those sampled (RSPv, MOs, and lateral (VISl) and primary (VISp) visual cortex). We ranked the cortical regions in the TD dataset by the fraction of low correlation pairs out of the total number of pairs per source (**Figure 5c**). Regions with the highest projection “specificity”, i.e., having more target-defined projection patterns that are different from the WT matches, included most of the prefrontal areas (ORBl, ACAv, ILA, prelimbic (PL), and medial orbital (ORBm)) and MOs. There was no relationship between the number of pairs tested in a source region and the fraction of low correlation pairs observed in that source (r=0.08, p=0.7). Notably, Spearman correlation coefficients were significantly lower for low correlation pairs in the prefrontal and medial modules compared with the visual module, and in the medial module compared with the prefrontal module (**Figure 5d,** p<0.02, one-way ANOVA followed by multiple t-tests).

We plotted the labeled cortical projections on surface views to visualize differences in the projection patterns of TD compared to WT experiments for low correlation pairs in one example source region from the prefrontal, visual, and medial modules (**Figure 5e-g**). As mentioned above, none of the TD experiments with ACAd as a source were a low correlation pair, and cortical projection patterns indeed looked similar on the surface view (see **Figure 5e** and **Figure S8**). In VISp, we found the correlation coefficients for 13% of all pairs were below the 95% prediction interval (10/80 pairs). One of these pairs was VISp neurons that target VISl (VISp_VISl_, n=3 replicates, **Figure 5f** and **Figure S9a**). VISp_VISl_ projections also target the laterointermediate (VISli) and and posteromedial (VISpm) visual areas but send few or no projections to anterolateral (VISal) and and rostrolateral (VISrl) visual areas. Notably, this pattern resembles a previously reported projection type based on L2/3 single cell axon mapping in VISp^61^ (*i.e.,* the LI-LM-PM projection type), indicating that the TD method can identify projection patterns similar to those seen in single neuron reconstruction experiments, and that at least some cells in L5 of VISp have projection patterns similar to the projection types previously identified in L2/3. Other examples of target-defined neuron classes that project to a subset of the WT projection pathway from VISp are provided in **Figure S9b-e**. In RSPv, TD experiments with the lowest r_s_ compared to their WT match included RSPv_PL_ projections which clearly represent a small portion of the WT pathway (**Figure 5g**). Specifically, most of these projections are to midline and other structures within the medial parts of the DMN mask, notably excluding much of the visual cortex outside of the DMN. These results show that some regions have more specificity in their target-defined projection patterns than others, and that for RSPv, some of these projection types could be related to the DMN.

### Two target-defined projection patterns in RSPv related to the DMN

RSPv is a core DMN region that is part of the posterior medial cortical network module. Relative to anterior DMN regions in the prefrontal module, we found that RSPv sends a more variable, but overall larger, fraction of projections to targets outside the DMN (**Figure 2i** and **Figure 4h**). RSPv was the only source in the medial module with low pairs, and the r_s_ values for its low correlation pairs were some of the smallest in the dataset (white points in **Figure 5c,** right panel). Thus, we focused next on determining in more detail how the projection patterns of the injection-matched experiments in RSPv relate to the DMN. We replicated the midline-projecting RSPv_PL_ pattern (**Figure 6a-c**), and identified a similar, slightly less restricted, pattern from RSPv_ACAd_ experiments (**Figure 6d-f**). In contrast, projections from RSPv_VISl/VISp_ experiments (targets in VISp or VISp and VISl) had a different portion of the WT pathway labeled, *i.e.,* a visual-projecting pattern (**Figure 6g-i**). These two projection types contact a distinct subset of WT targets (**Figure 6j,k**), with the midline-projecting RSPv_PL_ and RSPv_ACAd_ experiments primarily projecting to in-DMN targets and RSPv_VISp/VISl_ experiments contacting more out-DMN targets.

**Figure 6.**
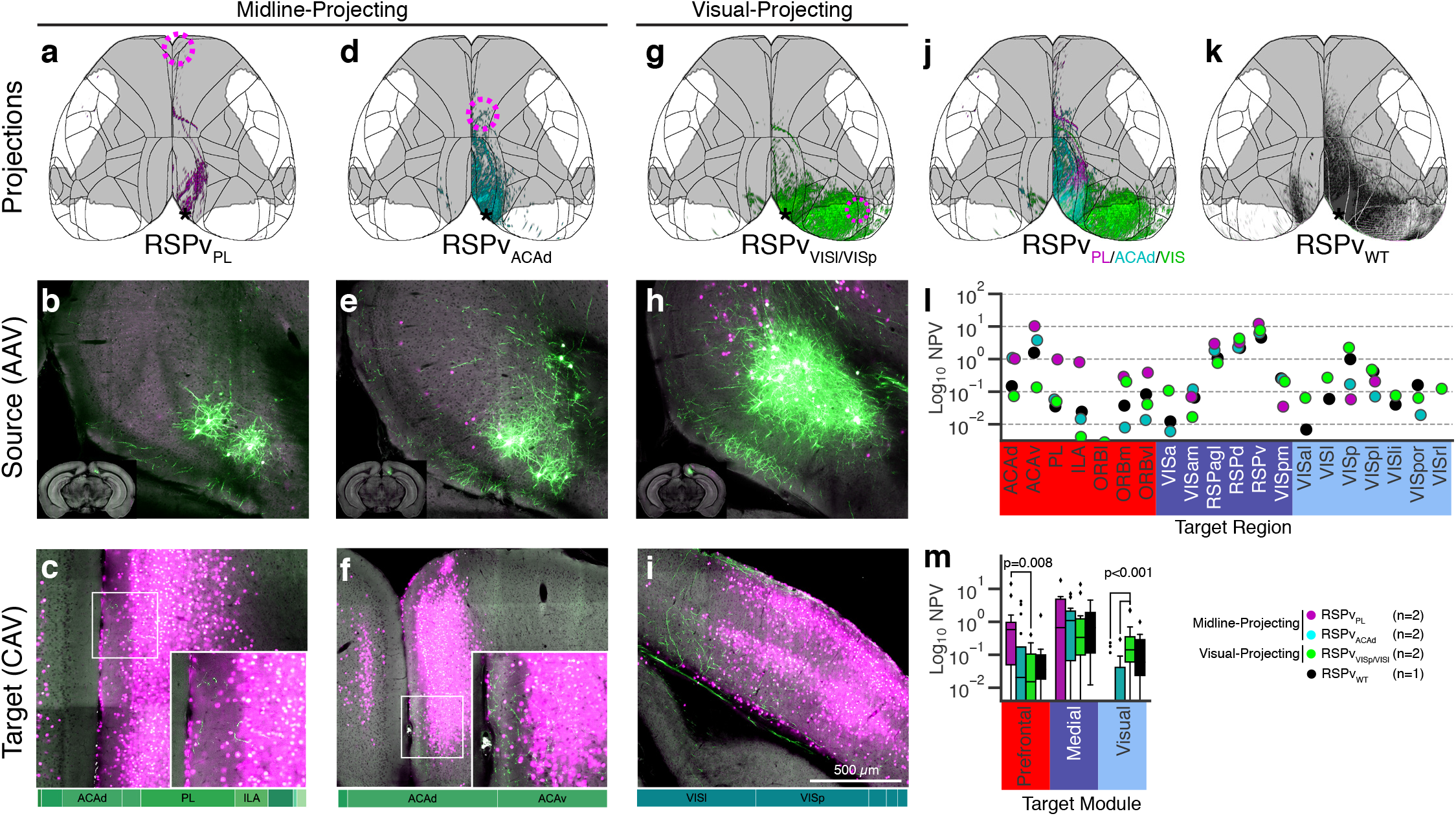
RSPv contains cells with in-DMN midline-projecting and out-DMN visual-projecting patterns. **a-i** Top-down cortical projections (**a,d,g**) and single image planes through the source (**b,e,h**) and target (**c,f,i**) injection sites for target-defined (TD) injection experiments in RSPv. Bars on the bottom of (**c,f,i**) show the regions in each CAV2-Cre injection site as the fraction of the injection polygon that overlapped with each region. PL and ACAd targets (**a-f**) have midline-projecting patterns, while VISl and VISp targets (**g-i**) have a visual-projecting pattern. **a-I** Full image series available online at http://connectivity.brain-map.org/projection/experiment/cortical_map/[insert experiment ID]. Experiment ids: RSPv_PL_ 592522663, RSPv_ACAd_ 521255975, RSPv_VISl/VISp_ 569904687, RSPv_WT_ 112595376. **j** overlay of the cortical projections from the three experiments shown in (**a,d,g**). **k** WT injection matched to the three TD injections in **j**. **l** Normalized projection volume (NPV) in selected targets plotted for each TD projection type (truncated at 10-2.5). **m** Projection strength to all targets in the Prefrontal, Medial, and Visual modules, thresholded at 10^−2.5^. Boxes=IQR, whiskers=1.5×IQR, outliers plotted as points, p-values were determined by multi-way ANOVA followed by Tukey’s post hoc test with correction for multiple comparisons.

We compared connection strengths from RSPv to selected ipsilateral cortical projection targets for the midline-projecting and visual-projecting RSPv cell populations (**Figure 6l**). Both types had axons in ACAd and ACAv, but connection strengths were higher for RSPv paired with the DMN vs. visual targets. We also compared the total projection strength from each RSPv TD projection type to all ipsilateral targets in the prefrontal, medial, and visual modules (**Figure 6m**). RSPv_PL_ experiments had stronger projections to prefrontal targets compared to RSPv_VIS_ experiments (p=0.008). Conversely, visual-projecting RSPv cells had stronger projections to targets in the visual module than either midline-projecting type (p<0.001, **Figure 6m**). Most higher visual areas received little to no input from the midline-projecting types, with the exception of the posterior-lateral visual area (VISpl), to which nearly all experiments projected (**Figure 6l**).

We checked whether midline DMN-projecting types were distributed in other RSPv subregions (**Figure S10a,b),** but found this pattern only in this ventral caudal location (analogous to A29a,b in the rat^66,67^). RSPv_ACAd_ projections from cells in the rostral or dorsal part of caudal RSPv had projection patterns that were more similar to their WT matches (r_s_ = 0.80±0.02, 0.88±0.06 for rostral and dorsal respectively). Specific midline-projecting cells were also labeled in an RSPv TD injection that involved VISpl and ENTm, but not in a similarly located TD experiment with CAV2-Cre centered in VISpl, but which did not include ENTm as a target (**Figure S10c,d**).

We noted that the midline-projecting RSPv injection experiments appeared to contain source cells located superficial to those from visual-projecting experiments (**Figure 6b,e,h**). Visual inspection of the images overlaid with each other suggests these cells are located in superficial L5 (**Figure 7a**). CCFv3 does not contain separate annotations for L2 and L3, and small variances in registration precision might result in the assignment of some voxels in deep L3 to L5 or vice versa (**Figure 7b**). Consistent with this, when we plotted the amount of signal in the injection site registered to layers in CCF space, we found that although most source cells were mapped to L5 for all experiments, the midline-projecting pattern had some signal assigned to L2/3, and visual-projecting experiments had some signal assigned to L6 (**Figure 7c**).Thus we further confirmed the relative sublayer location of these different projection types by comparing the distribution of source cells across the cortical depth with the distribution of tdTomato expression in four cortical layer-selective Cre driver lines crossed to the Ai14 reporter line, imaged with STPT and registered to CCFv3^52^; L2/3 (Cux2-CreERT2), L4/5 (Scnn1a-Tg3-Cre), L5 (Rbp4-Cre_KL100), and L6 (Ntsr1-Cre_GN220, projections in L5, **Figure 7d-g**). We used the cortical streamlines from CCFv3, which represent the shortest orthogonal path from the outer to inner cortical surface^52^, to plot the distribution of fluorescent intensity across cortical depth in RSPv (**Figure 7h**, left). We then compared these distributions to similar plots of eGFP fluorescence levels inside the injection site (**Figure 7h**, right). Notably, the peaks of the distributions for each of these three experiments are ~ 40% (midline-projecting) and 60% (visual-projecting) of the relative distance from the cortical surface, a range more like the L4/5 and L5 line Cre lines (Scnn1a and Rbp4) than either the L2/3 (Cux2) or L6 (Ntsr1) line. These plots further suggest that both projection types are indeed in L5, but midline-projecting neurons are located in a sublayer above most of the visually-projecting population.

**Figure 7.**
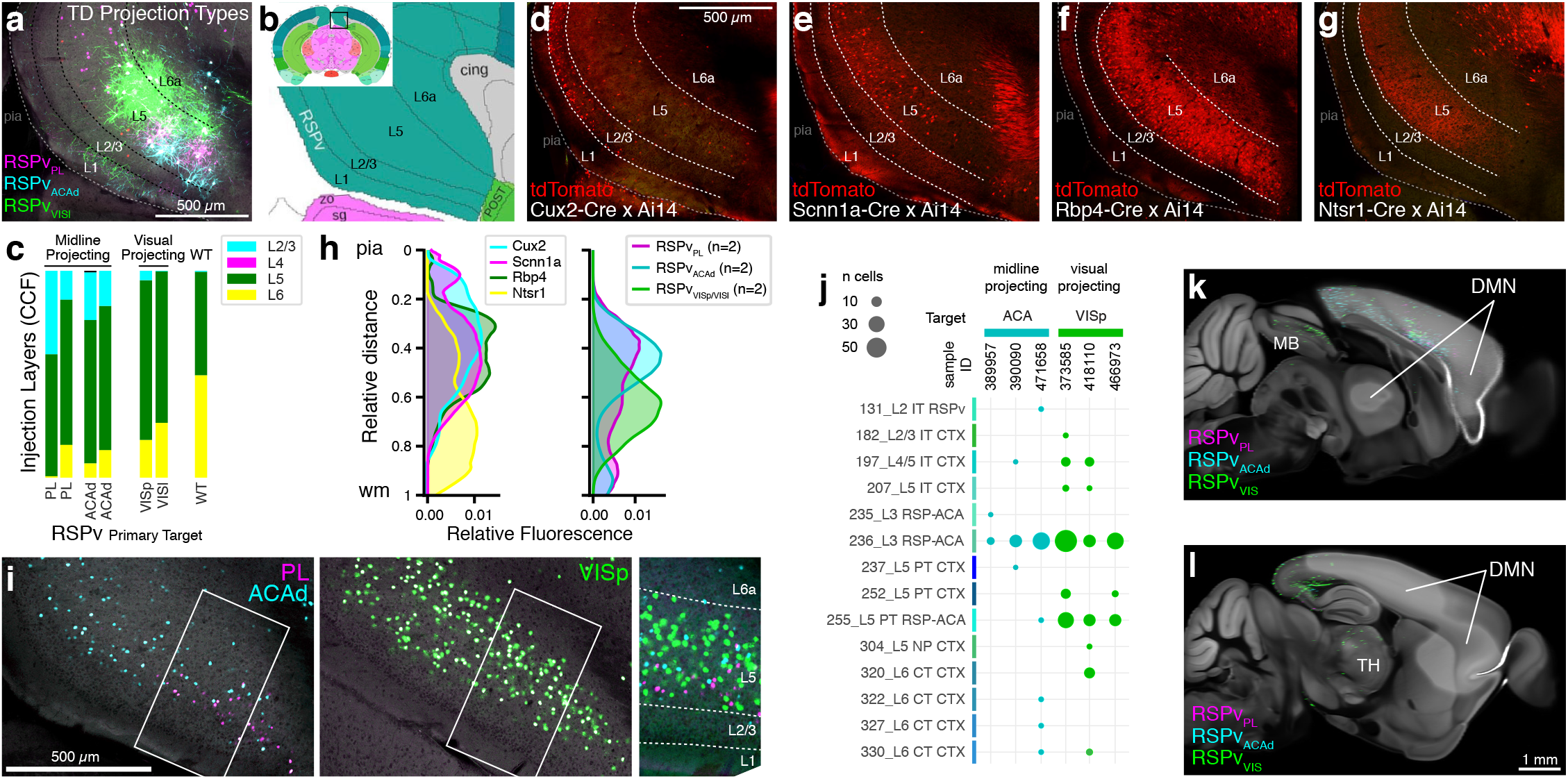
Midline-projecting RSPv cells are located in more superficial portion of layer 5 than visual-projecting cells, but they both map to the same unusual transcriptomic type. **a** Overlay of the single image planes from the TD source injection sites in **Figure 6 b,e,h**. Dashed lines indicate layer boundaries. **b** Coronal image from the Allen Reference Atlas showing the layer assignment for this portion of RSPv. **c** Layer composition of midline-projecting and visual-projecting TD source injections determined by registration to CCFv3. Midline-projecting experiments had more of their injection site assigned to L2/3 than visual-projecting experiments. **d-g** Single sections from layer-selective Cre transgenic mouse driver lines crossed to the Ai14 reporter line used to draw the layer boundaries in **a**. Note that tdTomato+ cell bodies for Ntsr1-Cre x Ai14 are in L6 but dense projections from these cells are also visible in L5 (**g**). **d-g** Full image series available online at http://connectivity.brain-map.org/static/referencedata/experiment/thumbnails/[insert experiment ID]?TRANSGENE=true&image_type=TWO_PHOTON, experiment IDs: Cux2 475359813, Scnn1a 475384080, Rbp4 475383272, Ntsr1 472986374. **h** Plots showing the fluorescence levels across depth from pia to white matter for tdTomato fluorescence in Cre driver lines crossed to the Ai14 reporter (left) and for EGFP fluorescence levels in TD experiments (right). **i** Single coronal planes in RSPv from two brains with retrograde trans-synaptic rabies tracer viruses injected in PL (magenta, left) and ACAd (cyan, left) or VISp (green, center). White box indicates region shown in a magnified view across cortical layers (right). **j** Retro-seq mapping of single-cell RNA sequence data collected from RSPv cells labeled with retrograde CAV2-Cre injections in ACA and VISp. Cluster numbers and labels from (Yao et al. 2020). **k,l** Sagittal sections from the CCFv3 mouse brain template at the midline (**k**) and slightly lateral to the midline (**l**). Segmented, gridded projections from the three experiments shown in **a** are overlaid. **k** Midbrain (MB) projections were observed in experiments with visual-projecting RSPvVISl/VISp cells (green) but not midline-projecting cells (cyan and magenta). **l** Projections in the thalamus (TH) were observed in experiments with visual-projecting cells but not midline-projecting cells.

Because of the known and previously described tropism of the CAV2-Cre virus for L5 neurons, we also performed replicate injections to label RSPv_ACAd_, RSPv_VISp_, and RSPv_VISpl_ projections, using the recently described nontoxic rabies variant RVΔGL-Cre which does not have this bias^65^. The separate types of midline-projecting RSPv_ACAd_ and RSPv_VISpl_ cells and visual-projecting RSPv_VIsp_ cells were also labeled with RVΔGL-Cre (**Figure S10e-g**). Critically, all experiments using both retrograde viruses had more L5 than L2/3 cells (**Figure S10h**), indicating that L2/3 cells are not contributing in a major way to these two projection types.

Finally, we also validated sublaminar differences of these projection types with data generated following retrograde transynaptic rabies tracing experiments. We injected rabies virus (CVS-N2CΔG) expressing nuclear-localized eGFP into PL, ACAd, or VISp^68^. RSPv L5 cells were labeled in all three cases, and the rabies-labeled cells providing input to PL were superficial to cells with projections to VISp. ACAd projecting cells were more intermingled with both PL and VISp projecting cells in these experiments (**Figure 7i**). We conclude that there are at least two projection types in RSPv L5, one that primarily projects to DMN targets (midline-projecting), and a second that primarily projects to visual targets (visual-projecting).

### Correspondence of midline- and visual-projection types with transcriptomically-defined cell types

Recently, we completed a large-scale study using single cell RNA sequencing (scRNA-seq) to derive a cell type taxonomy for the entire mouse isocortex and hippocampal formation^45^. Results showed that most transcriptomic cell types are common across the isocortex with some variation along spatial gradients, but a few regions do contain unique cell types, notably the midline structures RSP and ACA. So, we determined which transcriptomically-defined cell types contributed to the different RSPv projection-types by performing Retro-seq experiments^41^. We injected Cre reporter mice (Ai14, Ai75) with a retrograde Cre virus (CAV2 or rAAVretro^69^) in ACA or VISp, then isolated cells from the caudal part of RSPv by dissection followed by fluorescence-activated cell sorting (FACS). We obtained single cell transcriptomes using our SMART-Seq v4 (SSv4) pipeline^41^. Retro-seq cells were mapped to the clusters from the isocortex and hippocampal taxonomy^45^ by computing the correlation of gene expression for an individual cell with the median gene expression for each cell type within the taxonomy (**Figure 7j**). Most midline-projecting cells mapped to a single cluster, “236_L3 RSP-ACA”. Many of the visual-projecting cells also mapped to this cluster, but we found other transcriptomic types represented too, including “255_L5 PT RSP-ACA” for the visual-projecting cells (one ACA-projecting cell also mapped to this cluster). Thus, we did not find a 1:1 relationship between transcriptomic and RSPv projection type. The 236_L3 RSP-ACA cluster is an unusual subclass. It strongly expresses two L4 marker genes, *Scnn1a* and *Rspo1*, one of the L5 PT cell markers (*Bcl6,* but not *Fam84b*), and it also expresses a pan-IT marker, *Slc30a3*. Yao *et al.* concluded that this L3 RSP-ACA subclass has an intermediate cell type identity between IT and PT^45^. It was called “L3” because of mixed layer marker gene expression and because there is no anatomically-defined L4 in retrosplenial cortex. Our results above suggest that the location of these cells is more consistent with superficial L5. We also confirmed that whole brain projections labeled in the visual-projecting RSPv_VISp/VISl_ experiments are consistent with the “PT” classification; labeled axons reach to midbrain (**Figure 7k**) and thalamus (**Figure 7l**). In contrast, the midline-projecting types lacked subcortical projections, consistent with the “IT” cell class (**Figure 7k,l**). Thus, midline-projecting and visual-projecting patterns both seem to originate from cells in the “236_L3 RSP-ACA” cluster, which are likely a mix of IT and PT projection types as predicted by gene markers.

## Discussion

We used co-registered fMRI and mesoscale connectivity tracing with several viral and transgenic tools to identify the regional, layer, and cell-class connectivity of the mouse DMN. We found that DMN regions have strong preferential connectivity with each other, and that they overlap with two modules derived previously from analyses of structural connectivity (SC) network architecture^25^. We further found that L2/3 neurons in the DMN have primarily in-DMN projections while L5 neurons in DMN regions project to both in-DMN and out-DMN targets. Finally, we identified specific subclasses of in- and out-DMN projection neuron types in a core DMN region, retrosplenial cortex, and identified these cells based on their transcriptomic signature. Together, these findings provide an enhanced understanding of the specific cell classes and connection types that make up the functionally-defined DMN.

### Anatomical structures of the mouse DMN

By aligning our rsfMRI masks with the Allen CCFv3 reference atlas^52^, we comprehensively describe which regions brain-wide participate in the DMN. We report the overlap of the DMN with CCFv3 structures at 100 μm voxel resolution, which we used to produce a conservative list of DMN structures, all within the isocortex. The core DMN structures we identified using this method are in good agreement with previously published DMNs in rodents^3–7,34,70,71^. Exceptions included the hippocampus proper and retrohippocampal regions (including entorhinal cortex), which were not included in our DMN mask but were suggested to be to be part of the DMN in some studies^3,4,70^ but not others^5,6^. This inconsistency is likely to reflect differences in the DMN-detection strategies employed, which are strictly dependent on the number and quality of ICA components selected. To overcome this potential issue, in the present study we cross validated ICA-based DMN selection by comparing the resulting component topography with that obtained with seed-based probing of the anterior cingulate, highlighting a highly comparable configuration. Importantly, a very similar DMN topography was recently described using non-correlational rsfMRI dynamic mapping, leading to the identification of dynamic BOLD co-activation states closely recapitulating the antero-medial components of the DMN we mapped here^51^. Magnetic field inhomogeneity in the rodent ear canal, however, can limit the resolution of hippocampal and parahippocampal structures, ultimately impairing the reliable detection of more lateral areas in the rodent brain^3,72^. Importantly, recent work in humans has also posed a role for other subcortical structures in the DMN, including the thalamus^73^, a region that is robustly innervated by prefrontal DMN components^32^. Notably, we also found the DMN mask overlapped with voxels in the mouse medial thalamus, but to a much lesser extent than cortex. Since the DMN and core masks primarily overlapped with isocortex voxels, and all of our identified DMN regions were in the isocortex, we focused analyses on corticocortical structural connectivity. Future work could include mapping DMN connectivity from thalamus.

Our recently published modular structure of the mouse cortical network^25^ bore a striking resemblance to the DMN mask and to previously published modules derived from functional connectivity analyses^13,32^. The close alignment between the structures in these two modules and the DMN mask indicates that the DMN can be identified as a strongly connected network even against the background of all other cortical connections. Core DMN regions are in the prefrontal and medial modules, including orbital, anterior cingulate, prelimbic, retrosplenial, and parietal cortex. We found three DMN regions outside of the prefrontal and medial modules: secondary motor cortex and the trunk and lower limb regions of somatosensory cortex. The inclusion of MOs as a DMN structure was not surprising given its close spatial proximity to ACAd, and further examination of the difference in projection pattern for medial and lateral MOs revealed that the DMN does not precisely align with anatomical structure boundaries in that region. SSp-tr and SSp-ll, as primary sensory areas, were not expected to be DMN structures. However, a multi-site mapping of the mouse DMN at higher spatial resolution revealed a similar fringe involvement of primary sensory areas consistent with our findings here^7^, and Stafford et al. (2014) also identified portions of primary somatosensory cortex in their putative DMN.

The separation of the DMN into two modules is reminiscent of reports that the human DMN is composed of multiple interacting subnetworks^37,74,75^. However, direct comparisons are problematic since the rodent brain lacks clear homologs to some DMN regions such as the precuneus, posterior cingulate areas 23 and 31^67,76^ and the dorsolateral prefrontal cortex^77^. Some DMN regions appear to be conserved between primates and rodents, including the retrosplenial, anterior cingulate, and medial prefrontal cortex^3–6,70,77^. Recent cross-species probing of DMN dysfunction supports this notion, as comparable DMN-centered alterations in connectivity were recently observed in mouse and human subjects harboring autism-associated 16p11.2 chromosal microdeletions^78^.

### Organization of mouse DMN mesoscale connectivity

Across a variety of experiments and analyses, we consistently found that whether an injection site was inside or outside the DMN mask strongly predicted the fraction of its projections that were inside the DMN mask. This preferential connectivity inside the DMN agrees with previous reports of high correspondence between SC in anterograde tracing data from the Allen Mouse Connectivity Atlas (MCA) and DMN FC^6,34^. The expansion of the MCA to include projections from Cre-dependent tracer injections in layer-selective Cre driver lines^25^ enabled us to extend the analysis of preferential connectivity to cell classes defined by their layer of origin. Our finding that L2/3 IT cells in the DMN primarily project inside the DMN while L5 cells have both in- and out-DMN projections provides a new level of insight into the anatomical organization of the DMN. We also report the addition of 183 new TD viral tracing experiments, available through our data portal (http://connectivity.brain-map.org/), from which we used a subset here to investigate projection target specificity related to the DMN. We found evidence that cells projecting to one in-DMN target are more likely to project to other in-DMN targets, but only for sources in the medial module. Every source in the medial module that had TD experiments with both in- and out-DMN targets had a higher fraction of in-DMN projections when paired with an in-DMN target (**Figure 4h**). These results implicate medial module regions as potentially well-placed to mediate “switching” between different states.

Quantitative comparisons between viral tracer experiments are challenging because a large amount of the variation between experiments can be attributed to small variations in injection location ^79^. The smaller volume of our TD experiments posed an additional challenge for comparing them with larger injections in WT animals. We tackled this problem by rigorously quantifying the expected correlation between pairs of replicate injections in WT mice. The model we derived to predict the expected correlation between two experiments based on their injection volume, the distance between their centroids, and the overlap of their injection areas can be extended to future studies as a metric for quantifying the relative similarity between two experiments.

We also present evidence that some cortical regions have broadly distributed projection patterns, even at the level of target-defined neuron populations, whereas other regions contain cell populations with more specific, or restricted, target subsets. TD projections from most regions in the prefrontal module were particularly specific, but ACAd was a clear exception to this rule, with nearly all TD injections showing very broad projections like the matched WT experiments. This result is consistent with reports of ACAd being a key integrative hub region for the mouse DMN^32^. Recent studies have identified high diversity in the projection patterns of single neurons in auditory^80^, motor^81^, and visual^61^ cortex, indicating that cortical regions may generally have large variation in the specific targets contacted by individual cells. The existence of some regions with more stereotyped projection patterns would have implications for both the function of those regions (i.e. more integrative, hub-like) and for the number of single neuron reconstructions that will be required to identify all the projection types in these regions. Whether individual neurons within ACAd project to many or all of ACAd’s targets, or whether the specific set of targets we tested here (ORBvl, RSPv, RSPagl, VISam, MOp, and VISp) are shared among many projection types, may be better addressed through ongoing efforts to reconstruct the complete morphology of many single labeled neurons^82,83^.

We discovered a region in the posterior ventral part of RSPv that contained midline-projecting in-DMN cells and visual-projecting out-DMN cells. These projection types could be reproducibly separated by retrograde tracer injections in ACA, PL, and VISpl to label midline-projecting cells, or in VISp or VISl to label visual-projecting cells. The two projection classes could be distinguished based on their cortical projection targets, the presence or absence of subcortical projections (only visual-projecting cells had subcortical projections), and their depth from the pia surface. However, consistent with a recent report^84^, the two projection types could not be differentiated by their gene expression profile.

The midline-projecting RSPv cells were found in only one particular portion of RSPv, caudal to the corpus callosum splenium in the most ventral portion of RSPv, adjacent to postsubiculum. In rats, RSPv is parcellated into four regions, and midline-projecting RSPv cells were found in a region analogous to A29a,b, although this delineation does not currently exist in the Allen CCFv3. This region of RSPv was included in a medial subnetwork identified by Zingg et al. (2014), who noted its strong reciprocal connectivity with ACAv, ORBm, PL, and ILA, as well as its connections with medial entorhinal cortex and the subiculum. This network is particularly interesting with respect to the DMN, since the midline-projecting cells in RSPv are anatomically ideally situated to convey hippocampal output from the subiculum to medial prefrontal cortex. Alternating activation of midline-projecting and visual-projecting cells in RSPv could act as a switch between resting state DMN activity and visual processing in retrosplenial cortex. Functional characterization of the activity of these cells in various behavioral states would help clarify their potential role in the DMN.

### Relevance of mouse to human DMN comparisons

A unifying theory for cross-species comparisons of cortical DMN anatomy can be posed from analyses of spatial gradients in molecular, anatomical, and functional features across the primate cortex, which result in the hierarchical ordering of regions from primary sensory to transmodal association areas^56,85,86^. If the DMN indeed arises from temporally correlated activity between brain regions at the top of these hierarchies, its anatomical composition will differ between species based on the identity and location of polymodal, or association, cortex. Performing similar analyses of gradients in other species could therefore reveal both candidate DMN regions and conserved organizational principles^32,87,88^. Cortical hierarchies are also defined by laminar-based projection patterns into feedforward and feedback pathways^39^. In the mouse, we showed a shallow overall cortical hierarchy, but notably most of the DMN regions defined here using the functional DMN mask were at the top end of this purely anatomical-based hierarchy. Furthermore, when the six cortical modules were hierarchically ordered using laminar-based rules, the prefrontal and medial modules were at the top (along with the lateral module), while somatomotor, visual, and auditory modules were at the bottom^25^.

Our mesoscale data on cell class-specific connectivity also provides new testable predictions about the anatomical basis of functional networks. For example, L2/3 neurons may play a unique role in supporting DMN cohesion, cross-hemisphere synchronization of rs networks, and intra-module communication, while subclasses of L5 neurons may be involved in the coordination and switching across different brain-wide networks. The description of layer- and projection-specific cellular DMN components also provides potential inroads into understanding the selective vulnerability of the DMN to pathological processes in human diseases like Alzheimer’s^89^, autism, and schizophrenia^12^.

## Methods

### Mice

Experiments involving mice were approved by the Institutional Animal Care and Use Committees of the Allen Institute for Brain Science in accordance with NIH guidelines. Heterozygous Ai75 mice (RCL-nT; Jackson Laboratory #025106) on the C57Bl/6J background were used for all experiments except retro-seq, where homozygous Ai14 mice were also used (Jackson Laboratory # 007914). Tracer injections were performed in male and female mice age P57-P61, except for experiments in which the AAV injection was targeted by intrinsic signal imaging and the CAV2-Cre injection was coordinate-based (n=10 experiments). These 10 mice received a CAV2-Cre injection and implanted headpost at age P38-P42 and a second, ISI guided AAV-CAG_FLEX-EGFP injection at P53-P61. Injection ages for all mice are provided in **Supplementary Tables 3** and **4.** Mice were group-housed with a 12-h light-dark cycle. Food and water were provided ad libitum. rsfMRI was carried out on adult (12-18 week-old) male C57BI6 mice, purchased from the Jackson Laboratory (Bar Harbor, USA). Mice were group-housed with a 12-h light-dark cycle. Food and water were provided ad libitum.

### Functional MRI and DMN identification

Bold fMRI timeseries data were acquired from n=40 mice under light sedation (0.7% halothane, and artificial ventilation). rsfMRI acquisition parameters are described in greater detail in ^51^. Briefly, mice were anaesthetized with isoflurane (5% induction), intubated and artificially ventilated (2%, surgery). At the end of surgery, isoflurane was discontinued and substituted with halothane (0.75%). Functional data acquisition commenced 45 minutes after isoflurane cessation. Mean arterial blood pressure was recorded throughout imaging sessions. Arterial blood gases (p_a_CO_2_ and p_a_O2) were measured at the end of the functional time series to exclude non-physiological conditions.

All rsfMRI data were acquired with a 7.0 Tesla MRI scanner (Bruker Biospin, Ettlingen) using a 72 mm birdcage transmit coil, and a four-channel mouse brain for signal reception. Single-shot EPI time series were acquired using an echo planar imaging sequence with the following parameters: TR/TE 1200/15 ms, flip angle 30°, matrix 100 × 100, field of view 2 × 2 cm^2^, 18 coronal slices, slice thickness 0.50 mm, 500 volumes and a total rsfMRI acquisition time of 10 or 30 minutes, respectively.

To optimally register fMRI time-series to the 3D Allen Mouse Brain Common Coordinate Framework, v3 (CCFv3), we adopted the procedure recently described in ^90^. Briefly, fMRI time-series were first registered to an *in-house* T2-weighted mouse brain template with a spatial resolution of 0.1042 × 0.1042 × 0.5 mm^3^ (192 × 192 × 24 matrix^5^) using *fsl-flirt* and 12 degree-of-freedom. A low component group ICA analysis was then performed with GIFT toolbox (http://www.nitrc.org/projects/gift/) in the coordinate system of the template. Previous studies showed that the use of low component ICA (N = 5) permits to identify a DMN-like network in the mouse brain^5^. ICA analysis was run on demeaned data using the Infomax algorithm, with no autofill of data reduction values, and a brain mask to remove non-brain signal. All the other default parameters of GIFT (included PCA data reduction) were left unaltered. Independent components were scaled to z scores and thresholded at |Z| = 1, or |Z|=1.7 as in ^5^. These thresholds exceed thresholding levels employed in previous rodent rsfMRI studies^91–94^. To maximize component orthogonality and minimize the contribution of between-network anti-correlation^5,51^, only positive ICA weights were considered.

The results of DMN-based identification of ICA were independently corroborated by comparing the maps obtained with seed-based probing of the anterior cingulate, a core hub component of the rodent DMN^8^. To this purpose, a 3 × 3 seed voxel placed on the anterior cingulate of spatially-registered time series and Individual subject correlation maps were transformed to normally-distributed z scores using Fisher’s r-to-z transformation, before assessing group level connectivity distributions using one sample t-tests. Resulting group maps were thresholded at |T|= 8 (corresponding to uncorrected p< 0.00001, two-tailed test), followed by a cluster extent thresholding (p<0.01). The T threshold employed maximizes spatial correlation (Spearman rho 0.73) and dice coefficient (0.75) between |Z|= 1 ICA and seed-based maps.

### Projection of in-house mouse brain template into Allen CCFv3 Reference Space

We projected the in house MRI template (and co-registered ICA components) onto the CCFv3 using a combination of linear (affine) and non-linear (Syn) registrations using the ANTs package (*antsRegistration* command^95^. Before the registration, we initially aligned images used images intensity (-*r* option) to ensure an initial rough alignment. For both transformations, the optimisation was performed over five resolutions with a maximum of 100 iterations at the coarsest level. At the full resolution, 10 iterations were used for the affine registration, and 5 iterations were used for the non-linear registration. Smoothing and shrinking factors (7×5×5×3×1 and 9×7×5×3×1, respectively) were the same for both transformations. To enhance contrast, and hence help the registration, a slightly dilated brain mask was used. The gradient step was set to 0.1 for the affine registration, and to 0.15 for the Syn algorithm. In order to prevent non-realistic deformations, the spatial shifting in space for each iteration of the Syn algorithm was controlled by a smoothing factor (*updateFieldVarianceInVoxelSpace* = 3.0), whereas this was not the case for the total deformation (*totalFieldVarianceInVoxelSpace* = 0). For both the affine and the Syn algorithms, we adopted mutual information as similarity metric. The number of bins was set to 32, and values were sampled regularly in 50% of the voxels.

### Assignment of functional network masks to CCFv3 voxels

The parameters obtained from registering the bias field corrected in-house template were used to linearly project the ICA components into CCFv3 (*antsApplyTransforms*). The N3 algorithm as provided by ANTs was used to perform bias field correction (*N3BiasFieldCorrection:* Shrinking factor was set to 1, 50 iterations and 4 fitting levels were used). Registration quality was assessed by qualitatively comparing a brain mask mapped from the coordinate system of the in-house template into the CCFv3 reference space and the Allen “root” label (whole brain mask). Since the cerebellum (coronal slices 1 to 114) was a region of no interest, it was removed from the both images.

### Assignment of CCFv3 structures to fMRI masks

To determine which mouse brain structures were part of each ICA component in the fMRI dataset, we first computed the fraction of each fMRI map covered by each of the 12 structures in the CCFv3 ontology at the “coarse” level (structure set 2) plus “fiber tracts” by summing the number of voxels in the 100 μm resolution structure map that overlap with the fMRI mask and dividing by the total number of voxels in the fMRI mask. We further computed the overlap of the fMRI masks with each of the 316 “summary structures” from the Allen Reference Atlas (structure set 167587189) by summing the number of 100 μm voxels in each structure mask that overlapped with the fMRI mask and dividing by the total number of voxels in the structure. Selected structure overlaps with the DMN are presented in **Figure 1e**, and the overlap of all summary structures with the DMN and core masks are included in Supplementary Table 1. The code for computing the overlap of summary structures with any of the fMRI masks is available at [Github link to be added].

### Projection data analysis: DMN connectivity in WT experiments

We used 300 anterograde tracing experiments from the Allen Mouse Connectivity Atlas (MCA) with injections in the isocortex from WT C57Bl/6J mice (n=129), Emx1-IRES-Cre mice (n=66), and Rbp4-Cre_LK100 mice (n=105). We previously showed that projections in Emx1-Cre and Rbp4-Cre mice are highly correlated with experiments in WT mice^25^. This dataset can be viewed at http://connectivity.brain-map.org (Filter Source Structure(s): Isocortex, Filter Mouse Line: wild, Emx1-IRES-Cre, Rbp4-Cre_KL100). We used the 100μm grid volumes for this analysis: arrays of 100 μm voxels with values between 0 and 1 that indicate the density of segmented pixels inside each voxel for voxels inside the injection polygon (injection grid) and outside the injection polygon (projection grid) see^24^). To quantify the injection DMN fraction, we first divided the sum of the gridded injection voxels inside the DMN mask by the sum of all injection voxels. We restricted our projection analysis to cortical projections by applying the isocortex mask to the DMN mask, then computed the projection DMN fraction by summing the gridded projection voxels inside the isocortex-DMN mask and dividing by the total number of projection voxels within the isocortex mask. Left hemisphere injections were flipped to the right hemisphere for visualization purposes.

#### DMN Coefficient

To control for the expected higher projection density in voxels near the injection site, we calculated the distance between the injection centroid and all other 100 μm voxels in the isocortex. To eliminate false positive signal, we first thresholded each voxel at 1.6×10^−4^, which was the median signal per isocortex voxel in two experiments with failed AAV-FLEX-EGFP source injections (see ^96^ Figure S1 for more information about segmentation errors and false positive signal). We added an epsilon value of 0.01 to the projection density in each voxel, corresponding to the 95^th^ percentile of the false positive signal in the two experiments with failed AAV-FLEX-EGFP source injections. Each voxel was classified as in-DMN or out-DMN using the fMRI mask. We then used a general linear model to derive a “distance coefficient” (*β*_*dist*_) and a “DMN coefficient” (*β*_*dmn*_) for each experiment (equation 1).

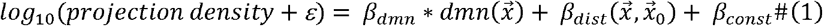

Where

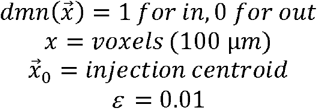

The correlation between the fraction of each injection that was inside the DMN and the DMN coefficient or the distance coefficient was calculated using ordinary least squares regression.

### Projection data analysis: Matched Cre Injections

We curated the set of injections from Extended Data Table 3^25^ so that all the injection experiments in each Rbp4 anchor group were either inside the DMN mask (injection DMN fraction > 0.5) or outside the DMN mask (injection DMN fraction < 0.5). In most sources, all experiments fell either inside or outside the DMN mask, but four source regions (MOs, SSp-bfd, VISpm, and VISp) each contained at least one experiment in both categories. We eliminated two out-DMN MOs injections (477037203 and 141602484) and two in-DMN VISp injections (300929973 and 272821309) and completely eliminated the VISpm group (Rbp4 anchor id = 485237081). Although barrel cortex (SSp-bfd) is not a DMN region, the matched group of injection experiments in SSp-bfd were mostly inside the DMN mask. We therefore also eliminated a single out-DMN SSp-bfd injection (178487444). One additional VISp experiment in a Rbp4-Cre mouse was manually classified as a PT projection type rather than an IT type, so that experiment was also dropped (517072832). This left 350 experiments in 24 sources (experiments from all six VISp anchor groups were combined into a single group). The distribution of experiments across DMN structures and layers was: L2/3: 18 in-DMN, 28 out-DMN; L4: 10 in-DMN, 33 out-DMN; L5: 26 in-DMN, 50 out-DMN; L5 PT: 35 in-DMN, 48 out-DMN; L6: 25 in-DMN, 36 out-DMN; all layers: 14 in-DMN, 27 out-DMN. Left hemisphere injections were flipped to the right hemisphere for visualization purposes.

### Viruses and injections

Atlas-derived stereotaxic coordinates were chosen for each source and target region based on The Mouse Brain in Stereotaxic Coordinates ^97^. Coordinates are reported as anterior-posterior (AP, referenced from bregma), medial-lateral (ML, distance from midline at bregma), and dorsal-ventral (DV, depth measured from the pial surface of the brain). Stereotaxic coordinates for 182 TD mesoscale connectivity mapping experiments are available in the online data portal in the experiment detail view. Stereotaxic coordinates for rv-ΔGL-Cre TD experiments, retrograde trans-synaptic tracing experiments, and retro-seq retrograde viral injection experiments are provided in **Supplementary Table 4**.

Many injections into visual areas were guided by sign maps derived from intrinsic signal imaging (ISI) through the skull as described in^25^; see also methods at http://help.brain-map.org/download/attachments/2818171/Connectivity_Overview.pdf. Sign maps for all ISI guided experiments are available in the online data portal as part of the cortical projection viewer. All viruses were injected into the left hemisphere, but images were flipped to the right hemisphere in figures to facilitate comparison with wild type experiments.

TD experiments used CAV2-Cre lot numbers 483, 738, and 990 with a titer of 1.0×10^12^ genome copies/ml (738) obtained from the Viral Vector Production Unit at the Universitat Autonoma de Barcelona, produced from a vector enabling expression of Cre under control of the CMV promoter^64^, paired with rAAV2/1-CAG-FLEX-EGFP-WPRE-bGH stock numbers V3900 and V5749 with a titer of 2.97×10^13^ and 1.34×10^13^ genome copies/ml (AAV-FLEX-EGFP) obtained from the Penn Vector Core. Three TD experiments (indicated in text) used RV-ΔGL-Cre at 4.26□×□10^10^ genome copies/ml as a retrograde tracer instead of CAV2-Cre, and two retro-seq experiments used AAV2-retro-EF1α-Cre at 5.0×10^12^ genome copies/ml as a retrograde tracer. rAAV and AAV2-retro were injected using iontophoretic current injection, 3 or 5 μA for 5 minutes at each depth specified in the stereotaxic coordinates or at 0.3 and 0.6 mm for ISI targeted injections. CAV2-Cre and and rv-ΔGL-Cre were injected via pressure using a Nanoject II system using 100-200 nl for each depth. Retrograde trans-synaptic input mapping experiments used the rAAV helper virus AAV1-DIO-TVA66T-dTom-N2cG with a titer between 3.60×10^12^ to 1.31×10^13^ genome copies/ml, produced from a vector enabling Cre-dependent expression of TVA receptor, rabies glycoprotein (G), and tdTomato in the cytoplasm of Cre-expressing infected neurons, paired with a G-deleted, ASLV type A (EnvA) pseudotyped rabies virus expressing a nuclear GFP reporter (RabV CVS-N2c(deltaG)-EGFP, Addgene vector ID 73461), with a titer of 5.00×10^9^ genome copies/ml injected 21 +/− 3 days after the initial rAAV helper virus injection^68,98^. Both viruses were produced at the Allen Institute^99^.

Twenty-one days after virus injection (9 days from the rabies virus injection date for retrograde trans-synaptic input mapping), mice were perfused with 4% paraformaldehyde (PFA), then brains were dissected and post-fixed in 4% PFA at room temperature for 3-6 hours, followed by overnight at 4ºC. Brains were stored in PBS with 0.1% sodium azide before proceeding to serial two photon (STPT) imaging using a TissueCyte 1000 system (TissueVision, Inc.) as described^24^. Briefly, coronal images were acquired every 100 μm through the entire rostral-caudal extent of the brain with xy resolution of 0.35 μm/pixel. Images shown in figures were cropped, downsampled, and dynamic range-adjusted using Adobe Photoshop CS6. To convert red images to magenta, the red channel was duplicated into the blue channel using Adobe Photoshop CS6. Left hemisphere injections were flipped to the right hemisphere for visualization purposes.

### Data Quality Control

There are 182 paired Cre-dependent anterograde AAV and retrograde CAV2-Cre tracer experiments available on the Allen Institute website (http://connectivity.brain-map.org filter by tracer type “Target EGFP”), of which 121 were included in this analysis. We excluded 11 experiments due to their source and target injection sites overlapping, and we only included experiments with cortical sources and targets. Twenty-six additional TD experiments with subcortical sources and twenty-five with subcortical targets are included in the online dataset but were not analyzed here. Replicate experiments were attempted for every source target pair, and 21 source-target pairs had n ≥ 2 experiments with the same primary source and target. One additional experiment has not yet been released online but is accessible through the AllenSDK (RSPv_VISp_ experiment # 868641659). One experiment (571410278) had 74% of its injection assigned to VISpl and 15% to VISpor, but its projection pattern was more similar to VISpor than VISpl so its primary injection structure was manually changed to VISpor for the purpose of identifying injection matched experiments. A list of all 183 TD experiments with their metadata and reason for inclusion/exclusion are included in **Supplementary Table 3**.

### CAV injection annotation and segmentation

Automated segmentation of EGFP signal was performed as previously described^57^ with minor modifications. When detecting EGFP-expressing fibers in the target-defined anterograde projection data, high intensity bleedthrough of nuclei in the CAV2-Cre injection site from the red channel into the green channel decreases the signal-to-noise ratio of the true fiber signal. We were unable to decrease the fluorescence intensity of the red channel because the autofluorescence from this channel is used for registration. Therefore, a supervised decision tree classifier was used to filter out segmentation artifacts based on morphological measurements, location, context, and the normalized intensities of all three channels. Regions in which this additional classifier were used were called “dense red zones”.

We evaluated the performance of the segmentation algorithm inside and outside of dense red zones by comparing segmented projections to manually traced projection data. We selected two image planes from three experiments that had regions of high red (ROI completely covered by dense red zone), no red (no dense red zone in ROI) and mid-red (ROI was partially covered by dense red zone). eGFP signal was manually traced in all ROIs by an expert annotator (JDW). The manual and automated segmentation were compared by calculating the Dice coefficient. Fewer of the manually traced axons were detected in the high red ROI than in the low-red ROI (p=0.01). For each experiment, the primary injection sites for the source (AAV) injection was determined as previously described^24^. The target (CAV2-Cre) injection site was determined using the boundaries of a second polygon that was manually drawn around the densest nuclear tdTomato labeling.

### Generation of a ground truth dataset for comparing replicate mesoscale viral tracing experiments

We previously used a distance threshold (≤ 500 μm) between injection centroids to identify spatially-matched anterograde tracing experiments^25^. However, this criterion alone was not sufficiently rigorous for TD – WT comparisons, in large part because TD injections had a smaller volume than WT injections (**Figure S6a**). The smallest TD injections had infected area diameters of 100-150 μm, so it was possible for the centroid of a TD source injection to be less than 500 μm from a WT experiment without the two experiments having any actual spatial overlap. Similarly, the centroid of a large WT injection could be located relatively far from a small TD injection, but the two experiments could still have large spatial overlap.

To determine the most important criteria for matching two injection experiments, we started by manually checking 1568 TD – WT experiment pairs and rating their match quality by visual inspection using the cortical projection view (available for any experiment at http://connectivity.brain-map.org/projection/experiment/cortical_map/[experiment_id]) and individual image sections in the injection site for both experiments (ratings provided in **Supplementary Table 5**). These ratings were used to choose thresholds for the percent of the injection volume contained in the primary injection structure, the distance between centroids, and the overlap between injection volumes based on Dice Coefficient that included most well-matched injection experiments and excluded most poor matches. We then applied these thresholds to pairs of WT experiments matched by injection structure to identify 627 pairs of experiments in the isocortex and that we considered replicates. The criteria for calling two experiments an injection match were:

1. The two experiments had the same primary injection structure (structure containing the highest fraction of the injection density, see ^24^).
2. If the primary injection structure made up less than 60% of the injection volume in either experiment, the secondary injection structure was also the same in both experiments (n=69 pairs in this category).
3. The two experiments had injection centroids located < 800 μm apart.
4. The Dice Coefficient was greater than 0.05.

Right hemisphere injections were flipped to the left hemisphere to calculate overlap and distance. Experiments meeting these criteria were called “injection-matched” to distinguish them from groups of experiments with the same source injection structure that were not necessarily spatially matched.

This group of “WT” experiments consisted of injections of anterograde AAV-EGFP tracer in WT C57Bl/6 mice (n=88 experiments), a Cre-dependent EGFP tracer (AAV-FLEX-EGFP) in Emx1-IRES-Cre and Rbp4-Cre_KL100 mice (Emx1: n=26, Rbp4: n=54 experiments), and a Cre-dependent synaptophysin-EGFP fusion protein tracer (AAV-FLEX-SypEGFP) in Emx1 and Rbp4 mice (Emx1: n=27, Rbp4: n=13 experiments) for a total of 208 experiments. This dataset contained at least one pair of injections in 27 of the 43 cortical regions. An additional seven cortical regions were represented as matched secondary sources in experiments with <60% of the injection volume in the primary structure.

We included Cre-dependent tracer injection experiments done in Emx1 and Rbp4 transgenic mice in the dataset we call WT replicates because the projection patterns from cortical injection experiments in these two mouse lines have been previously shown to be highly correlated with injections of anterograde EGFP tracer in WT mice^25^. However, we confirmed that there were no differences in correlations between experiments performed in WT mice and in these two Cre lines for this dataset. We calculated the Spearman correlation (r_s_) for each pair of experiments using the normalized projection volume (NPV) in ipsilateral and contralateral cortical target structures. To minimize the contribution of false positives, NPV values were thresholded at log10(NPV) = −1.5 before calculating correlation coefficients^25^.

The Spearman correlation did not differ for the three viruses (EGFP, Cre-dependent EGFP, and Cre-dependent SypEGFP), but transgenic line was a significant factor (p=0.13 for virus, p<0.0001 for transgenic line, two-way ANOVA). Post-hoc testing showed that no transgenic line pairs had r_s_ values significantly lower than WT – WT pairs. However, r_s_ values for Emx1 – Emx1 experiment pairs were significantly higher than several other pairs of transgenic lines (Emx1 – Emx1 vs. WT – WT: p=0.02, WT – Rbp4: p=0.001, Emx1 – Rbp4: p=0.008). In addition, WT – Emx1 correlations were significantly higher than WT – Rbp4 correlations (p=0.006). We did not exclude any of these pairs from the replicate dataset based on these statistics. For the full set of WT replicates used to construct the correlation model (all Cre line/virus combinations), r_s_ was 0.86±0.07.

### Correlation model selection

Even after applying our injection match criteria, injection volume, distance between centroids, and injection overlap were all significant factors that predicted the magnitude of the Spearman correlation (p<0.001 for all, multi-way ANOVA). We fit a simple model that predicts the r_s_ between two experiments given these three factors using the Dice coefficient as a metric of overlap and residual sum of squares (RSS) as the optimality criterion. We tested several simple functions relating r_s_ to these factors. The relationship between the smaller injection volume and the Spearman was better fit by an exponential function compared to logarithmic (RSS = 2.92 logarithmic vs. 2.80 exponential). The relationships between the centroid distance and Dice coefficient and r_s_ were both linear (RSS for distance = 3.10 linear, 3.18 exponential; RSS for Dice coefficient = 2.99 linear and 2.99 exponential). We started with the exponential model based on injection volume and tested whether it was improved by the addition of parameters for the Dice coefficient or distance. We derived p-values for the additional parameters by performing an F-test on the sum of squared residuals for the two models. The addition of parameters for distance or Dice coefficient further improved the model (RSS with distance parameter = 2.73, p=0.0001 vs. reduced model, RSS with Dice coefficient parameter = 2.71, p=5e^−6^ vs. reduced model). Including both distance and Dice coefficient further improved the fit (RSS=2.68, p=0.0005 vs. model based on size and distance). The final model is presented in equation 3 and **Figure S6d**.

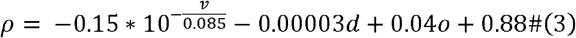

Where

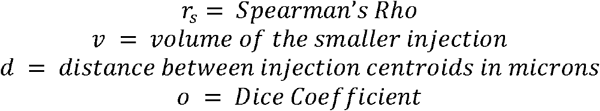

### Selection of wild-type matches for target-defined experiments

Using the same injection match criteria as we used for constructing the control dataset, we identified 586 pairs of matched TD and WT experiments, including at least one matched WT for all except 11 TD injections. The WT experiments matches consisted of injections of anterograde EGFP tracer (rAAV2/1- hSyn-EGFP-WPRE-bGH) in WT C57Bl/6 mice (n=49 experiments), Cre-dependent EGFP tracer (rAAV2/1-CAG-FLEX-EGFP-WPRE-bG) in Emx1-IRES-Cre and Rbp4-Cre_KL100 mice (n=18 and n=32 experiments), and a Cre-dependent synaptophysin-EGFP fusion protein tracer (rAAV2/1.pCAG.FLEX.synaptophysinEGFP.WPRE.bGH) in Emx1-IRES-Cre and Rbp4-Cre_KL100 mice (n=19 and n=10 experiments). 108 of 128 WT experiments matched to TD injections had also been used to fit the model.

### Identifying matched experiment sets

We used the same criteria to identify the largest groups of injection matched TD experiments that contained at least one experiment with an in-DMN target and one with an out-DMN target. Using the injection match criteria, we also identified all the WT experiments that matched with every TD experiment in each set, allowing one mismatch for each WT experiment (i.e. a WT experiment could match with every TD experiment in a group except one and still be included in a “set”). The experiment ids in each of the ten injection matched experiment groups are presented in **Supplementary Table 3.**

### Single-cell RNA-sequencing

Cells were collected by microdissection of RSPv from brains of mice with retrograde label. Single-cell suspensions were created and cells were collected using fluorescence activated cell sorting (FACS). FACS gates were selective for cells with tdTomato-positive label. Single cells were sorted into individual wells of 8-well PCR strips containing lysis buffer from the SMART-Seq v4 kit with RNase inhibitor (0.17 U/μl), immediately frozen on dry ice, and stored at −80°C. Cells were later processed for single cell RNA-seq using SMART-Seq v4 Ultra Low Input RNA Kit for Sequencing (Takara Cat# 634894) as previously described^41,45^.

Sequence alignment was performed using STAR v2.5.3^100^ in the two-pass mode. PCR duplicates were masked and removed using STAR option ‘bamRemoveDuplicates’. Only uniquely aligned reads were used for gene quantification. To identify their cell types, sequenced cells were mapped to the mouse cortex and hippocampus taxonomy^45^. The median gene expression for each cell type within the taxonomy was calculated using the SMART-Seq v4 dataset as reference. Cells were mapped by computing the correlation of gene expression of an individual cell with the median gene expression for each cell type. To estimate the robustness of mapping, mapping was repeated 100 times, each time using 80% of randomly sampled genes, and the probability for each cell to map to every reference cluster was computed. The cell type with the highest probability of mapping was chosen as the corresponding cell type of that cell.

### Statistics

Unless otherwise indicated, ranges of values are reported as mean ± SD. Correlation coefficients reported as “r” refer to the Pearson correlation, and as “r_s_” refer to the Spearman correlation. Unless otherwise indicated, p-values were computed using multi-way ANOVA with type II sum-of-squares and Tukey’s post hoc test for factors that were significant in the ANOVA. Significance was established using a p-value of 0.05. When multiple t-tests were performed for post-hoc analyses, we used the Benjamini/Hochberg correction for multiple comparisons.

### Data accessibility

Raw fMRI timeseries are available at these links (Part #1 of the dataset, N=20 mice, https://doi.org/10.17632/7y6xr753g4.1; Part #2, N=20 mice, https://doi.org/10.17632/r2w865c959.1) fMRI masks can be downloaded in nearly raw raster data (nrrd) format at [URL to be added]. Additional software for accessing the TD dataset and reproducing the analysis and figures in this publication is available at [URL to be added].

## Supporting information

Supplementary Table 1 Summary Structures DMN Overlap

Supplementary Table 2 DMN Structures

Supplementary Table 3 Target-Defined Dataset

Supplementary Table 4 Experiments Not Online

Supplementary Table 5 Injection-Matched Pairs

## Acknowledgements

The authors would like to thank Joaquín Goñi for helpful discussion on the analysis of layer-specific connectivity in the DMN. We thank Kiet Ngo, Mike Taormina, and Nadezhda Dotson for performing STPT imaging; Andy Cho, Jennifer Luviano, Linzy Casal, and Robert Howard for viral injections and intrinsic signal imaging; the Animal Care, Transgenic Colony Management, and Laboratory Animal Services teams for mouse husbandry and tissue preparation; and the Program Management team for administrative support. The project described was supported by the National Institute on Aging and the National Institutes of Health by Award Number R01AG047589 to J.A.H. Its contents are solely the responsibility of the authors and does not necessarily represent the official views of the National Institutes of Health. We thank the Allen Institute founder, Paul G. Allen, for his vision, encouragement, and support. This work was also supported in part by the European Research Council (ERC, DISCONN to A.G., Grant Agreement 802371). A.G. also acknowledges funding by the Simons Foundation (SFARI 400101, A. Gozzi), the Brain and Behavior Foundation (2017 NARSAD, Independent Investigator Grant 25861), the NIH (1R21MH116473-01A1), Telethon Foundation (GGP19177) and the University of Padua inter-departmental project Proactive.

## Author contributions

Conceptualization: J.A.H., J.D.W., A.G., H.Z. Supervision: J.A.H., A.G., S.M., H.Z., L.N., P.A.G., J.D.W., Project administration: W.W. Investigation, validation, methodology and formal analyses: J.D.W., A.L., K.E.H., P.B., A.W., P.A.G., N.G., J.E.K., A.H., P.R.N., T.N.N., E.G., C.T.J.V., O.F., M.N., A.M.H., N.D., K.A.S., B.L., B.T., S.M., A.G., J.A.H. Data curation: J.D.W., A.L., L.C., K.E.H., T.N.N., J.A.H. Software: N.G., L.K., J.E.K., W.W., D.F. Visualization: J.D.W., A.L., L.C., K.E.H., W.W., T.N.N., D.F., M.N., A.G. The original draft was written by J.D.W and J.A.H., with input from A.G. All co-authors reviewed the manuscript. The authors declare no competing interests. Correspondence and requests for materials should be addressed to J.A.H.

---

**Figure S1.**
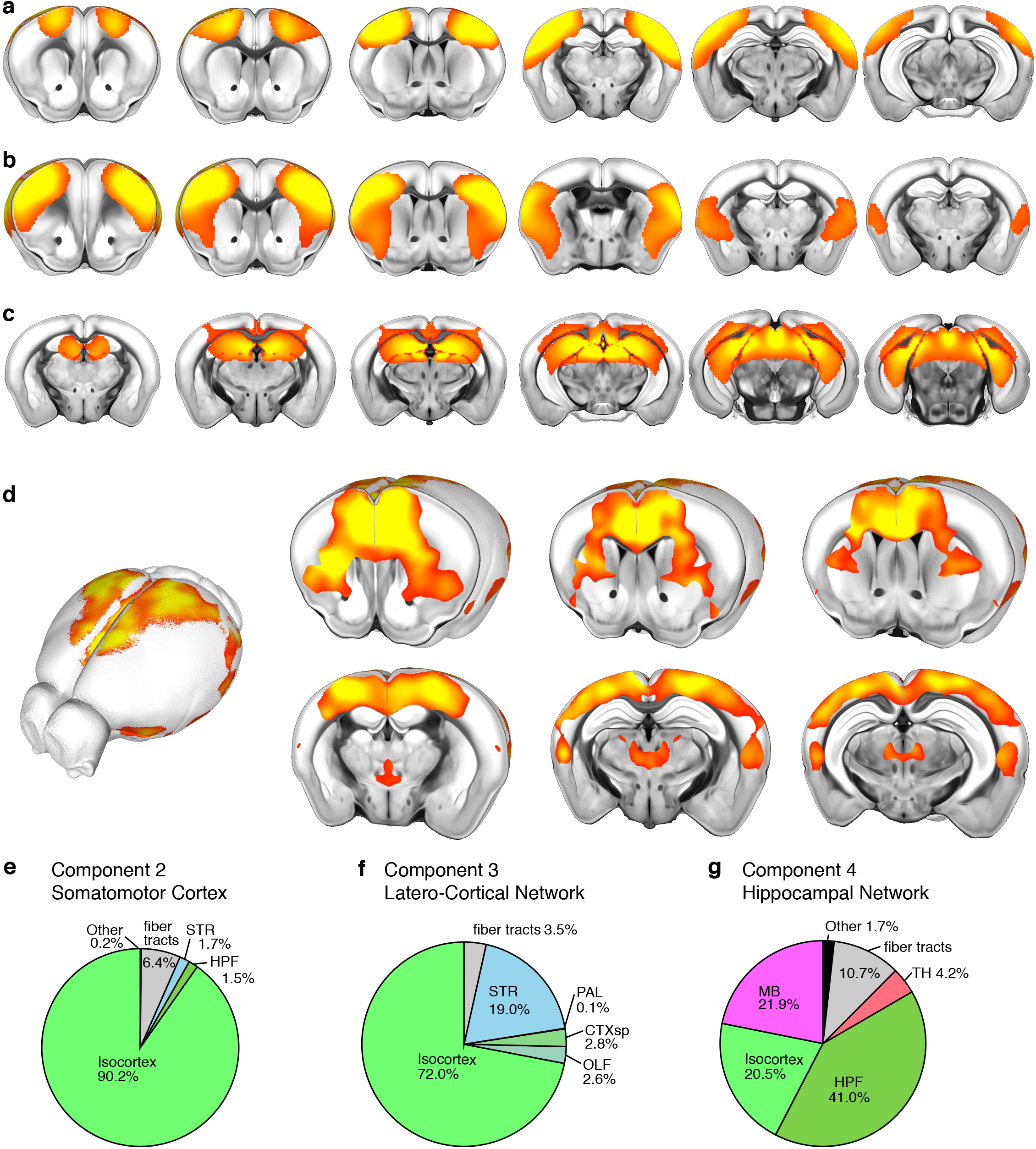
Overlap of functional networks with major brain divisions. **a-c** Additional components identified with ICA analysis on fMRI data. **d** An alternative DMN component generated with a seed pixel in ACA (Cg) resembles the ICA map in **Figure 1b,c**. **e-g** Pie charts showing the composition of each component in terms of major brain divisions. Abbreviations: TH: thalamus, MB: midbrain, OLF: olfactory areas, CTXsp: cortical subplate, HPF: hippocampal formation, STR: striatum, PAL: pallidum.

**Figure S2.**
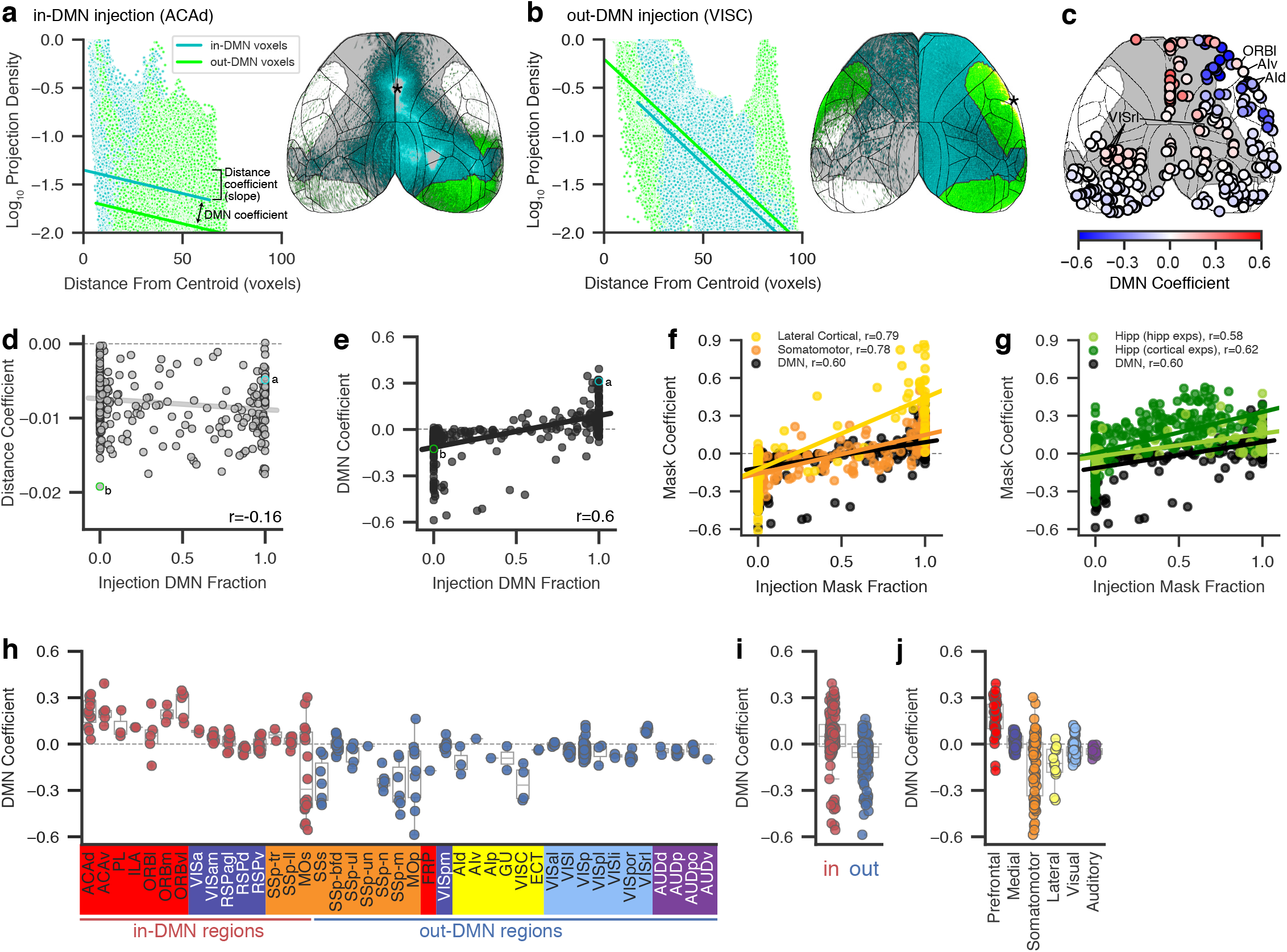
Preferential DMN connectivity independent of distance. **a,b** Projection density per voxel in isocortex plotted for in-DMN voxels (cyan points) and out-DMN voxels (green points) in one in-DMN experiment (**a**) and one out-DMN experiment (**b**). Lines show the model fit for in-DMN voxels (cyan line) and out-DMN voxels (green line). The magnitude and sign of the DMN coefficient is reflected in the distance between the in-DMN and out-DMN fit lines, and the magnitude of the distance coefficient by the slope of the lines. Abbreviations: ACAd, anterior cingulate area, dorsal part; VISC, visceral area. Experiment IDs: 125833030 (**a**), 180436360 (**b**). **c** Top down cortical projection map showing the location of these experiments colored by their DMN coefficient. Abbreviations: ORBl, orbital area, lateral part; VISrl, rostrolateral visual area; AId and AIv, agranular insular area, dorsal and ventral parts, respectively. **d** distance coefficient and **e** DMN coefficient for each of the 300 injection experiments in WT, Emx1-Cre, and Rbp4-Cre mice from the Allen Mouse Connectivity Atlas (MCA). r refers to the Pearson correlation. Experiments shown in **a** and **b** are labeled on the graphs in **c** and **d** (green and cyan borders). **f,g** Coefficients for the projections of cortical injection experiments in the lateral cortical and somatomotor ICA networks (**f**) and hippocampal network (**g**) as a function of the fraction of the injection inside the ICA mask for that network. The plot for the DMN coefficient from **e** is replicated for reference. The coefficient for 53 hippocampal experiments from the MCA is also shown for the hippocampal network (**g**, light green). **h** The DMN coefficient for each of the experiments shown in **c** grouped by their injection source. Sources are arranged by module and DMN membership. Boxplots show the range of values for all experiments in each source, with individual points for each experiment. **i** The experiments in **c** grouped by in-DMN and out-DMN regions. **j** The experiments in **c** grouped by module.

**Figure S3.**
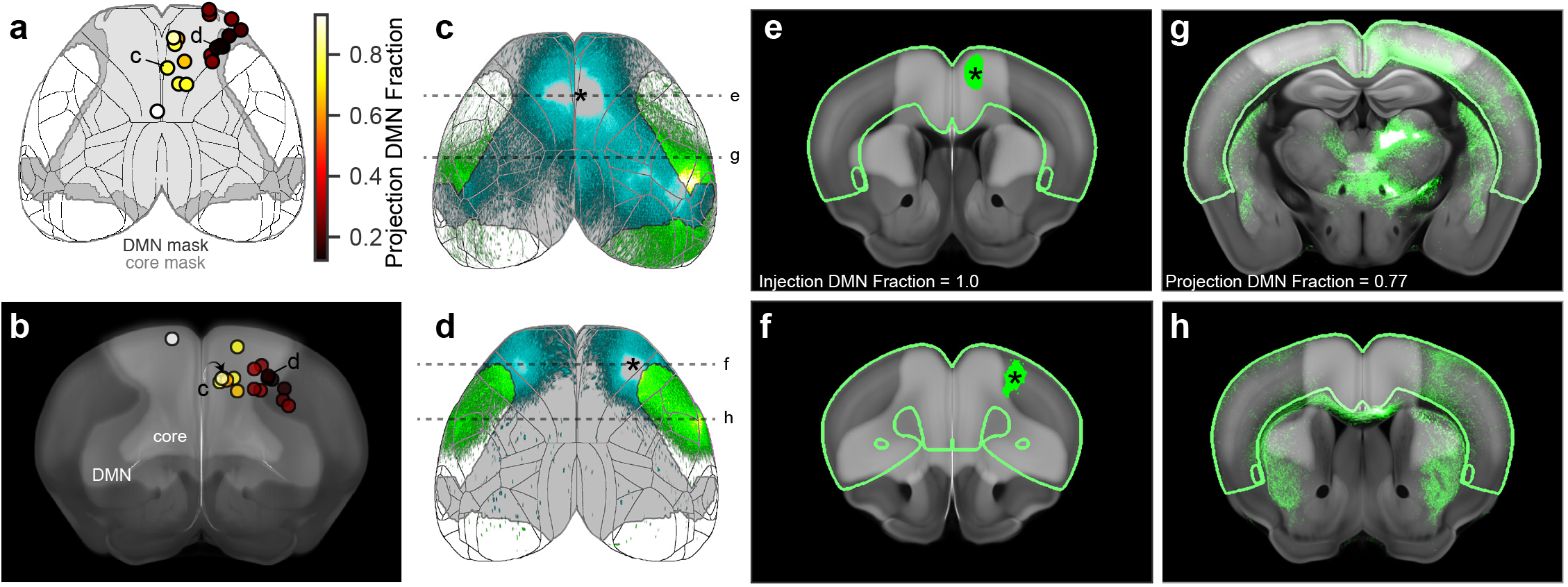
Secondary motor cortex has in-DMN and out-DMN subregions. **a** Cortical surface map showing the location of the 18 injection experiments in secondary motor cortex (MOs) in WT, Emx1-Cre, and Rbp4-Cre mice from the Allen Mouse Connectivity Atlas. The boundary of the DMN and core masks are shown in gray and cortical structure boundaries in black. Colormap indicates the percent of cortical projections inside the DMN mask for each experiment. Experiments shown in **c** and **d** are labeled. **b** Coronal view of the injection experiments in **a** overlaid on a maximum intensity projection of the corresponding portion of the average template brain. The DMN and core masks overlaid in light gray. Experiments are colored as in **a**. Experiments shown in **c** and **d** are labeled. **c,d** Cortical projection images showing two examples of MOs injections. Asterisks indicate the approximate centroid of the injections. In-DMN projections are pseudocolored cyan and out-DMN projections green. Full image series available online at http://connectivity.brain-map.org/projection/experiment/cortical_map/[insert experiment ID]. Experiment IDs: 141603190 (c), 595025284 (d). Dashed lines show the rostral-caudal position of panels **e-h**. **e,f** Single coronal image section through the center of the injection sites showing segmented injection pixels (green) overlaid on the corresponding coronal section from the CCFv3 template and DMN and core masks (light gray). **g,h** Single coronal sections at the approximate rostral-caudal position with the highest projection density showing segmented projection pixels (green) overlaid on the corresponding CCFv3 template section and DMN and core masks (light gray).

**Figure S4.**
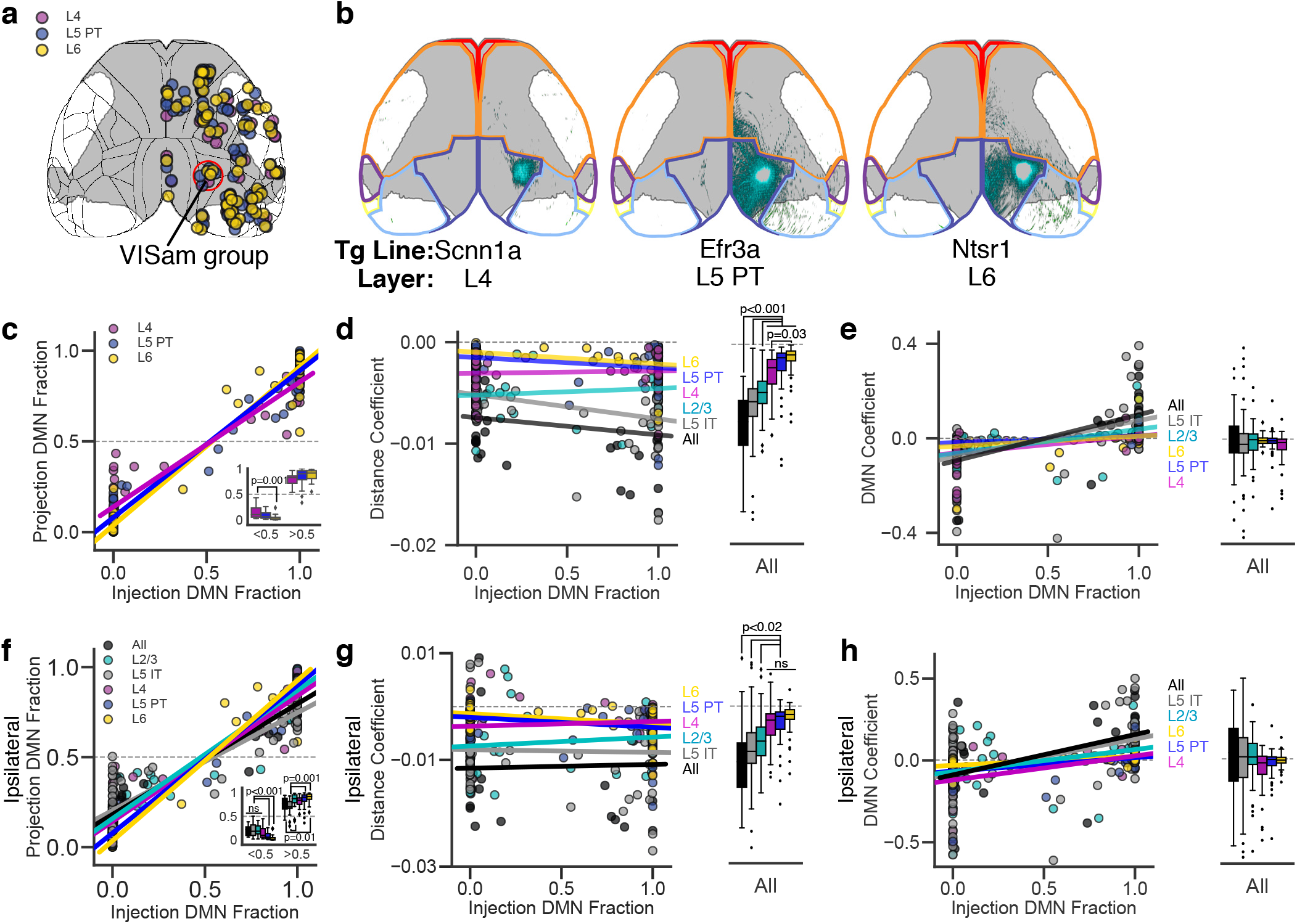
L4, L6. and L5 PT neurons have mainly local projections. **a** Location of matched injection experiments in L4 selective Cre lines (magenta, n=43), L5 PT selective Cre lines (blue, n=83), and L6 selective Cre driver lines (yellow, n=61). Red circles indicate the injection centroids belonging to the anteromedial visual cortex (VISam) group, shown in **b**. Gray overlay shows the DMN mask borders. **b** Top down cortical projections of experiments from the VISam group in a Scnn1a mouse line selective for L4 neurons, an Efr3a mouse line selective for L5 PT neurons, and an Ntsr1 mouse line selective for L6 neurons. Full image series available online at http://connectivity.brain-map.org/projection/experiment/cortical_map/[insert experiment ID]. Experiment IDs: Scnn1a 268038969, Efr3a 309515141, Ntsr1 156671933. **c** Scatterplot and linear fit for the percent of cortical projections inside the DMN as a function of the percent of the injection inside the DMN for each of the 187 cortical injections shown in **a**. Inset shows the fraction of DMN projections for injections binned by the fraction of the injection polygon inside the DMN mask. **d** Distance coefficient, and **e** DMN coefficient plotted as a function of injection DMN fraction for the set of layer-selective experiments shown in Figure 3a and S4a. Boxplots show the same points binned by the fraction of injection polygon inside the DMN mask (boxes=IQR, whiskers=1.5×IQR). **f** The fraction of ipsilateral projections inside the DMN as a function of the fraction of the injection inside the DMN. Inset shows the fraction of DMN projections for injections binned by the fraction of the injection polygon inside the DMN mask. The relative values for the different layer selective Cre lines are not different from the fraction of ipsilateral and contralateral projections inside the DMN (compare with **c** and **Figure 3c**). **g** The relationship between the distance coefficient and the fraction of an injection that is inside the DMN. The relative order of the Cre lines is the same as for ipsilateral and contralateral projections (**d**). **h** The relationship between the DMN coefficient for ipsilateral projections and the fraction of an injection experiment that is inside the DMN. The relative order of the Cre lines is the same as for ipsilateral and contralateral projections (**e**).

**Figure S5.**
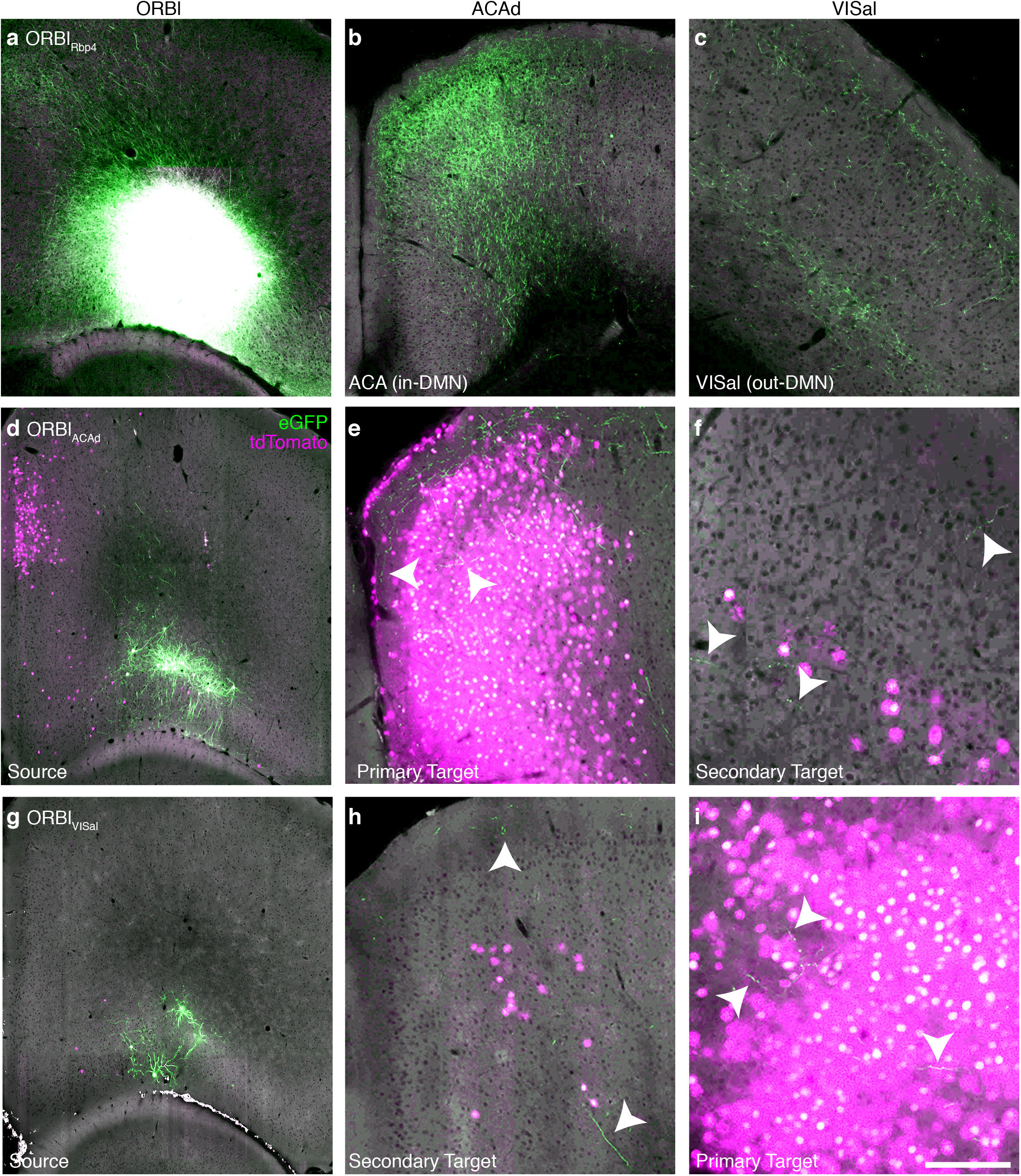
High magnification images of TD injection experiments. **a** One coronal image section through the center of an ORBl injection in an Rbp4-Cre mouse. The injection site is saturated in this image so that axons can be resolved. Projections from this experiment are visible in ACAd (**b**) and VISal (**c**). **d** One section through the approximate center of an ORBl_ACAd_ source injection (in-DMN target) showing cells coinfected with both viruses (green), and nuclei from cells infected with only CAV2-Cre (magenta). Projections from the ORBl_ACAd_ cells are shown inside the CAV2-Cre injection site (**e**, primary target, arrowheads) and in VISal (**f**, secondary target). **g** An ORBl_VISal_ source injection (out-DMN target). **h** Projections from ORBl_VISal_ cells in ACAd, which is a secondary target for this experiment. **i** Projections from ORBl_VISal_ cells in the primary target region, VISal (arrowheads). Scale = 500 μm. Full image series available online at http://connectivity.brain-map.org/projection/experiment/cortical_map/[insert experiment ID]. Experiment IDs: ORBl_Rbp4_ 156741826, ORBl_ACAd_ 571816813, ORBl_VISal_ 601804603.

**Figure S6.**
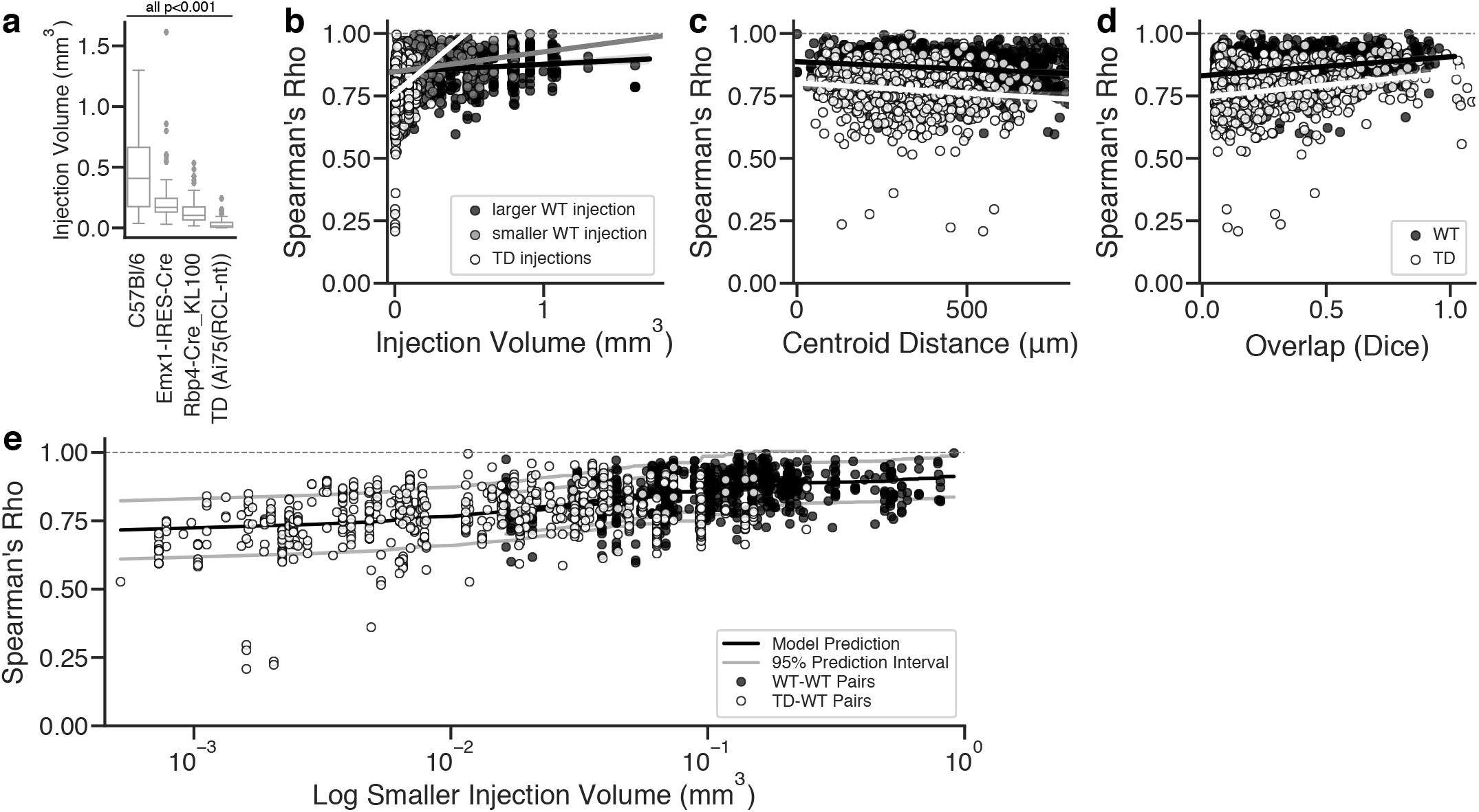
Comparing TD experiments with their WT matches. **a** Injection volumes in TD experiments are significantly smaller than injection volumes in WT, Emx1-Cre, and Rbp4-Cre experiments. Injection volumes are calculated from segmented pixels inside a polygon manually drawn around infected cells^24^. **b-d** Injection volume (**b**), distance between injection centroids (**c**), and Dice overlap (**d**) for WT-WT experiment pairs (black and gray) and TD experiments (white). **e** Predicted (black line) and actual (points) Spearman correlation for each WT-WT (black), and WT-TD (white) injection-matched pair. Gray lines show the 95% prediction interval.

**Figure S7.**
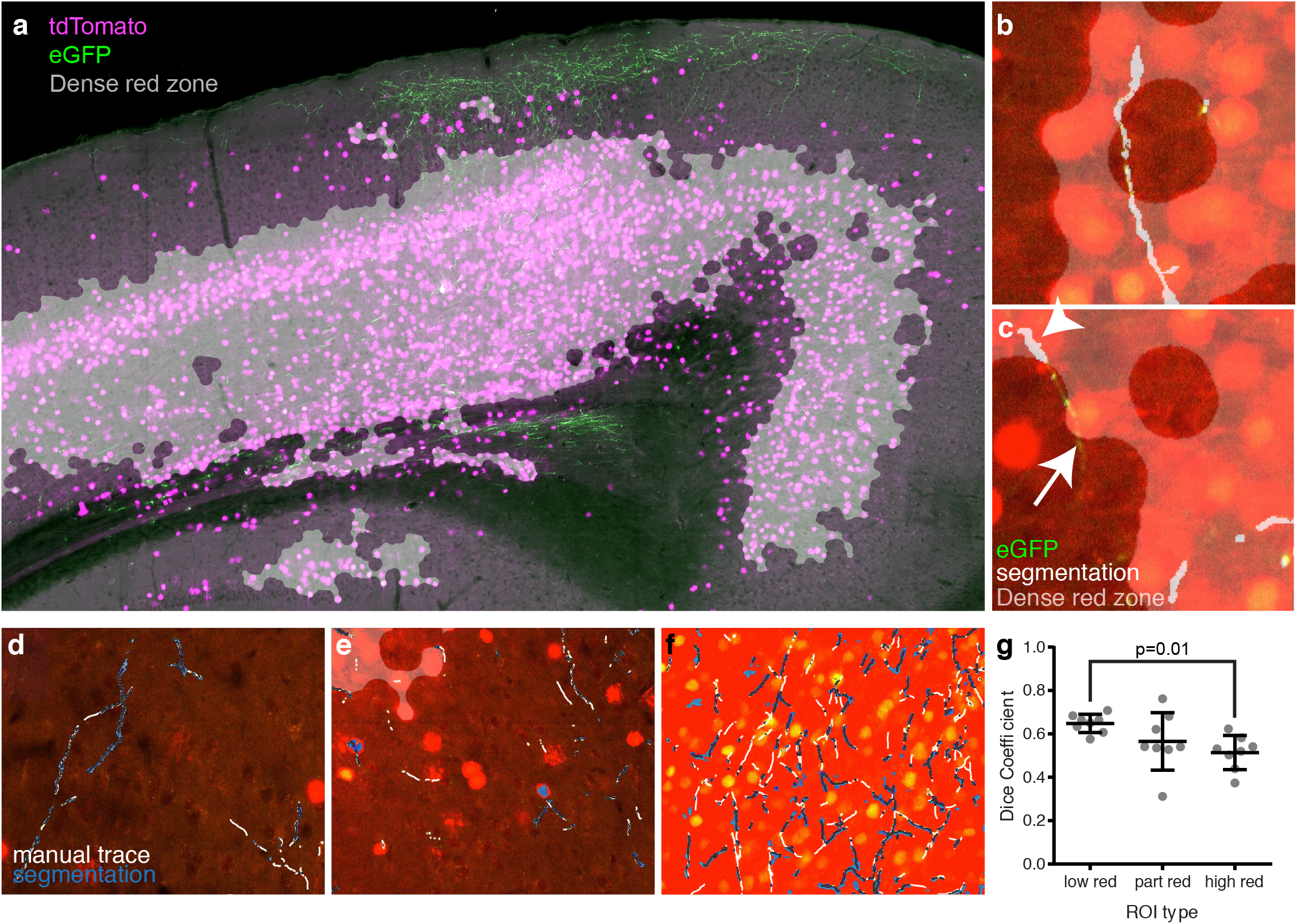
Performance of segmentation algorithm. **a** Image from a target-defined projection experiment showing retrograde Cre-dependent tdTomato expression in nuclei and cytosolic Cre-dependent eGFP expression in projecting fibers. The “dense red zone” identified by our segmentation algorithm is overlaid in gray. **b,c** eGFP+ axons crossing the border of the dense red zone (overlaid in gray) and the segmentation result for these fibers (white). In some cases, axons were detected both inside and outside the dense red zone (**b**), but other axons were detected inside (arrowhead) but not outside (arrow) the dense red zone (**c**). **d-f** Examples of manually traced axons in low red (**d**), mid-red (**e**), and high-red (**f**) regions. **g** The dice coefficient for manually traced and segmented axons was lower in high red regions than in low red regions.

**Figure S8.**
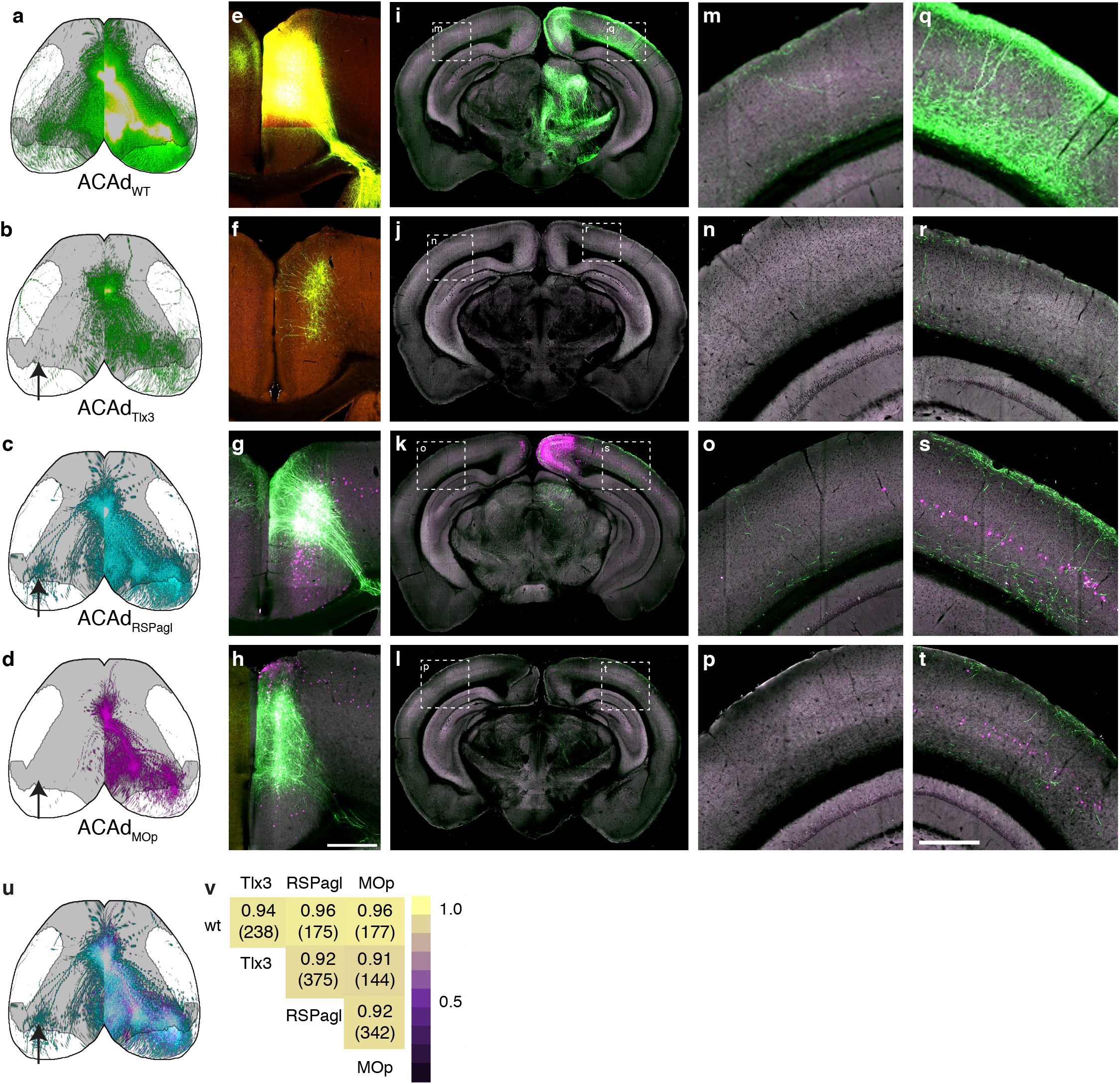
ACAd target-defined projections are homogeneous. **a-d** Top down cortical projection views for an ACAd injection in a WT mouse (**a**), a L5-selective Cre driver mouse (Tlx3, **b**), and two TD experiments (**c,d**). **e-h**. Single section image at the approximate center of each injection site. **i-l**. One coronal section from each experiment. Boxed insets show the area enlarged in **m-p** (contralateral) and **q-t** (ipsilateral). **u** Overlay of the top down cortical projections from the two TD experiments in **c** and **d**. Arrows in **b,c,d,u** indicate the only notable difference between these experiments: the presence of stronger contralateral projections in the ACAd_RSPagl_ experiment. **v** Correlation matrix for the four ACAd experiments shown here. Values indicate the Spearman correlation between each pair and the distance between the two injection centroids (in parentheses). Scale = 500 *μ*m. Full image series available online at http://connectivity.brain-map.org/projection/experiment/cortical_map/[insert experiment ID]. Experiment IDs: ACAd_WT_ 146593590, ACAd_Tlx3_ 293432575, ACAd_RSPagl_ 607059419, ACAd_MOp_ 475829896.

**Figure S9.**
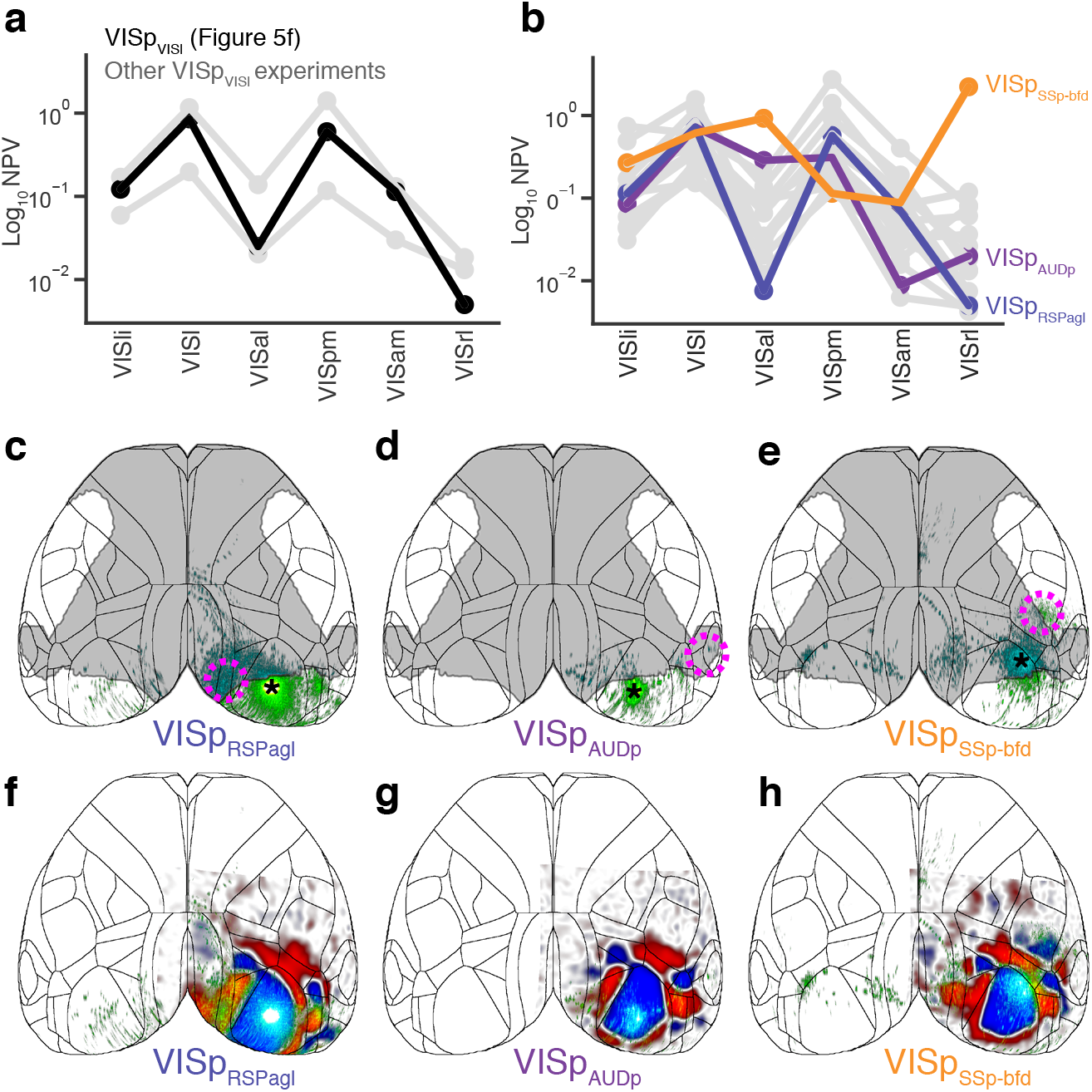
Different target-defined projection patterns from VISp sources. **a** Normalized Projection Volume (NPV) in six visual and medial structures for three VISp_VISl_ TD experiments (gray), and for the single VISp_VISl_ experiment shown in Figure 5f (black). **b** NPV in six visual and medial structures for 16 TD injections in VISp (gray). The VISp_RSPagl_ projection pattern (blue) resembles the VISp_VISl_ pattern in panel **a**, while the VISp_AUDp_ (purple) projections had a different pattern with strong projections in VISli, VISl, VISal, and VISpm and no projections to VISam and VISrl. VISp_SSp-bfd_ (orange) projections had yet another pattern, with projections to all the visual areas included here, and strongest in VISrl and VISal. **c-e** Top down cortical projection views of the VISp_RSPagl_, VISp_AUDp_, and VISp_SSp-bfd_ experiments shown in **b**. Asterisk indicates the approximate location of the AAV source injection centroid and dashed circles indicate the approximate area of the CAV2-Cre target injection. Full image series available online at http://connectivity.brain-map.org/projection/experiment/cortical_map/[insert experiment ID]. Experiment IDs: VISp_RSPagl_ 501787135, VISp_AUDp_ 501837158, VISp_SSp-bfd_ 539323512. The sign maps for the three example VISp TD experiments shown in **b** are overlaid on the top-down cortical projection views in **f-h**. The location of the injection centroid for the VISp_SSp-bfd_ injection relative to the sign map shows that this source injection was located in a different part of VISp than the VISp_VISl_ and VISp_RSPagl_ injections, more rostral and lateral. These subtle differences in source location and their correlation to different projection patterns emphasize the importance of careful experiment matching for quantitative comparisons between TD experiments.

**Figure S10.**
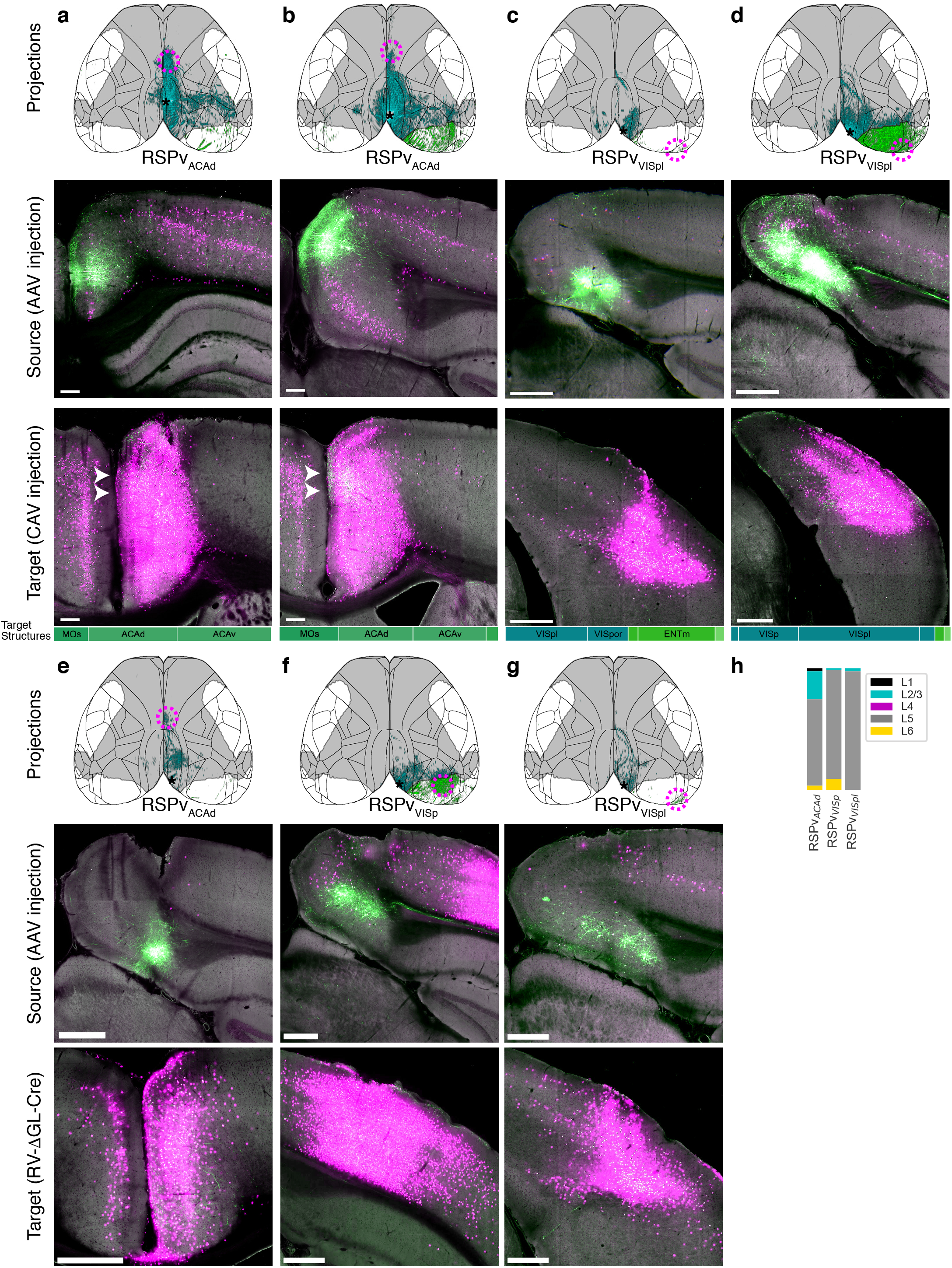
Midline-projecting RSPv_ACAd_ cells are only found in caudal, ventral parts of retrosplenial cortex, and can be labeled with either CAV2-Cre or RV-ΔGL-Cre target injections. **a-d** TD experiments with CAV2-Cre virus target injections. *top*: top-down cortical projections. Asterisk indicates the approximate centroid of the source injection and magenta circles indicate the approximate location of the CAV2-Cre target injection. *middle*: Single image section from the approximate center of the source injection site showing the location of EGFP-expressing cell bodies for cells that were infected with both viruses. *bottom*: Single image through the approximate center of the target injection site. Bars on the bottom of the images show the regions in each CAV2-Cre injection site as the fraction of the injection polygon that overlapped with each region. **a** An RSPv_ACAd_ TD injection with a source in the rostral portion of RSPv does not have a midline-projecting pattern. **b** an RSPv_ACAd_ TD injection with a source in the dorsal portion of caudal RSPv does not have a midline-projecting pattern. Arrowheads in **a,b** point to more dense axons in ACAd than ACAv. **c** Midline-projecting RSPv cells can be labeled with a target CAV2-Cre injection in VISpl. **d** A different RSPv_VISpl_ TD experiment did not label midline-projecting cells. **a-d** Full image series available online at http://connectivity.brain-map.org/projection/experiment/cortical_map/[insert experiment ID]. Experiment IDs: RSPv_ACAd_ (a, rostral) 475830603, RSPv_ACAd_ (b, dorsal) 571647261, RSPv_VISpl/ENTm_ (c) 592724077, RSPv_VISpl_ (d) 666090944. **e-g** TD experiments with RV-ΔGL-Cre target injections. *top*: top-down cortical projections. Asterisk indicates the approximate centroid of the source injection and magenta circles indicate the approximate location of the RV-ΔGL-Cre target injection. *middle*: Single image section from the approximate center of the source injection site showing the location of EGFP-expressing cell bodies for cells that were infected with both viruses. *bottom*: Single image through the approximate center of the target injection site. **e** A RSPv_ACAd_ TD experiment with a midline-projecting pattern. **f** A RSPv_VISp_ experiment with a visual-projecting pattern. **g** A RSPv_VISpl_ experiment with a midline-projecting pattern. **e-g** Experimental data available through the AllenSDK. Experiment IDs: RSPv_ACAd_ 605112318, RSPv_VISp_ 595890081, RSPv_VISpl_ 595261714. **h** Layer composition of each of the three TD source injections with RV-ΔGL-Cre as the target virus determined by registration to CCFv3. Scale = 500 μm.

## Notes

http://connectivity.brain-map.org/

https://doi.org/10.17632/7y6xr753g4.1

https://doi.org/10.17632/r2w865c959.1

### Competing Interest Statement

The authors have declared no competing interest.

